# Paradoxical gene regulation explained by competition for genomic sites

**DOI:** 10.1101/2025.11.27.691022

**Authors:** Krishna Manoj Aravind, Dhruv D. Jatkar, M. Ali Al-Radhawi, Eduardo D. Sontag, Domitilla Del Vecchio

## Abstract

Understanding how opposing regulatory factors shape gene expression is essential for understanding complex biological systems. A motivating observation, drawn from cancer epigenetics, is that removing an activating factor can sometimes lead to higher, not lower, expression of a gene that is also subject to a repressing factor. Prior theoretical work explained this counterintuitive behavior by competition of repressors and activators for genomic binding sites. However, it has been difficult to test this directly in natural systems, where layers of regulation obscure causal relationships. This paper introduces a fully synthetic, tunable genetic platform in a prokaryotic model system that reconstitutes this competition mechanism in a controlled and isolated setting. The genetic platform contains a target gene with binding sites for both an activator and a repressor, together with separate overlapping decoy binding sites for the same regulators. Activator and repressor functions are implemented using CRISPRa and CRISPRi, which permit independent control of regulator expression levels, design of the binding sites, and modulation of the binding affinities. Using this minimal system, we demonstrate that increasing activator expression level can reduce expression of the target gene when both regulators are present, consistent with the hypothesis that additional activator molecules displace the repressor from decoy sites, which becomes available to repress the target. By demonstrating how competition for genomic binding sites can invert expected regulatory responses, this synthetic framework provides a system for understanding similar paradoxical behaviors in natural regulatory networks and establishes a foundation for future studies in more complex mammalian contexts.

**Significance Statement:** Gene regulation is often described in terms of activators that increase expression and repressors that decrease it, yet biological systems frequently display counterintuitive behaviors. Here we show that competition between regulators for shared genomic binding sites can invert expected responses, so that increasing an activator can reduce target gene expression. Using a minimal, fully controllable synthetic system based on CRISPR activation and interference, we isolate and experimentally validate this mechanism. Our results demonstrate that such paradoxical effects arise not from changes in intrinsic regulatory roles but from redistribution of regulators across competing sites. This work provides a general, mechanistic framework for understanding nonintuitive gene-expression patterns observed in complex systems, including those relevant to disease.

## Introduction

Gene regulation is often described in terms of activating and repressing interactions, yet many biological systems exhibit responses that deviate from simple monotonic expectations. Increasing the abundance of a transcriptional activator does not always increase expression of its targets and, in certain contexts, can even reduce transcriptional output. These paradoxical effects have motivated efforts to better understand the mechanisms through which TFs shape regulatory behavior. Two complementary perspectives have emerged for understanding how TFs generate activating or repressing regulatory outcomes. One line of work focuses on the *intrinsic* regulatory role of a TF: the local mechanisms by which a transcription factor modulates RNA polymerase (RNAP) activity at a promoter. In this view, a TF may act “dually,” functioning either as an activator or as a repressor by favoring or disfavoring RNAP binding or initiation. Such duality can arise from the presence of both activation and repression domains within the same TF (1). More generally, the effective sign and strength of regulation depend on molecular context, including TF–RNAP interaction strength (2), promoter strength (3), TF–DNA binding affinity (4), promoter geometry, and binding location. This intrinsic perspective seeks to isolate promoter-level behavior by removing network-level confounders such as competing binding sites, feedback loops, and physiological coupling. Regulation is then described in terms of local TF– RNAP interactions, often decomposed into mechanistic contributions such as stabilization or destabilization of polymerase binding and acceleration or deceleration of transcriptional initiation (2). Rather than classifying TFs categorically as activators or repressors, this framework emphasizes a continuous spectrum of regulatory modes. TF concentration primarily moves the system along the response curve associated with a given molecular context, while the qualitative sign of regulation is determined by the underlying TF–RNAP kinetic mechanisms (2, 4).

A second, complementary perspective emphasizes the *extrinsic* context in which regulation occurs. In natural systems, TFs operate within shared pools of cofactors, polymerases, and genomic binding sites, and regulatory outcomes can depend strongly on resource allocation and competition (5–7). Classical examples include transcriptional squelching and decoy-site competition, in which increasing TF abundance redistributes limiting molecular resources and can produce counterintuitive responses such as paradoxical repression or nonmonotonic input–output curves (5, 7, 8). In this systems-level view, TF concentration does not merely scale a fixed regulatory curve; instead, it reshapes the effective regulatory landscape by altering the availability of active TF or cofactors across the network, so that apparent switches between activation and repression may arise from global constraints rather than from intrinsic properties of the TF at a single promoter (6, 7).

Consistent with this extrinsic perspective, early studies of transcriptional squelching demonstrated that overexpression of an activator can inhibit transcription by titrating limiting cofactors, highlighting that regulatory roles depend on network context rather than solely on intrinsic properties of individual proteins (5). A genetically engineered approach to ameliorating the squelching effect has been proposed, modeled, and implemented in mammalian cells (9). A central organizing principle underlying these effects is retroactivity, whereby downstream binding interactions feed back onto upstream regulator availability (10–13). When multiple regulators share binding resources, perturbations that alter binding elsewhere in the genome can redistribute transcription factors and reshape gene expression patterns. Decoy binding sites provide a clear illustration of this mechanism: both synthetic and natural systems have shown that additional binding sites can sequester regulators, modify effective concentrations, and alter regulatory dose response curves without altering the intrinsic sign of regulation (6, 7, 14).

These intrinsic and extrinsic viewpoints should therefore be regarded as complementary layers of understanding rather than competing explanations. Intrinsic models provide a local, mechanistic characterization of how a TF modifies transcriptional kinetics at a bound promoter, whereas resource-competition frameworks capture system-level effects that emerge when many targets and molecular pools interact simultaneously. A unified picture suggests that observed gene-expression responses reflect the interplay between these levels: local TF–RNAP mechanisms define the baseline regulatory mode, while network context determines how strongly that mode is expressed and whether it may appear inverted or attenuated in vivo (2, 6, 7, 13).

The intrinsic and extrinsic mechanisms discussed above are often formulated in terms of a single TF acting on a target promoter. However, gene regulation in vivo typically involves multiple regulators whose activities are coupled through shared binding pools and limited resources. To study this far more realistic scenario, especially for systems with both activators and repressors, raises fundamentally new questions. How does competition between distinct TF species reshape regulatory input–output relationships? Motivated by this perspective, we combine theoretical modeling with a controllable synthetic genetic platform to study the minimal principles underlying competition-driven regulation. We developed a mechanistic reaction network model in which an activator and a repressor regulate a target gene while also binding to decoy sites that represent the broader genomic environment (15). The model predicts that when activator and repressor molecules compete for overlapping decoy sites, increasing the activator concentration can release repressor molecules from those sites and thereby enhance repression at the target. This mechanism generates a nonintuitive input–output curve in which induction of an activator leads to reduced target expression, as illustrated in Fig. 1. Guided by these predictions, we constructed a minimal circuit in bacterial cells using CRISPR activation and CRISPR interference as synthetic activator and repressor modules. Overlapping binding sequences enforce competitive binding at decoy sites, and binding affinities are tuned through engineered mismatches at the target locus.

**Fig. 1.**
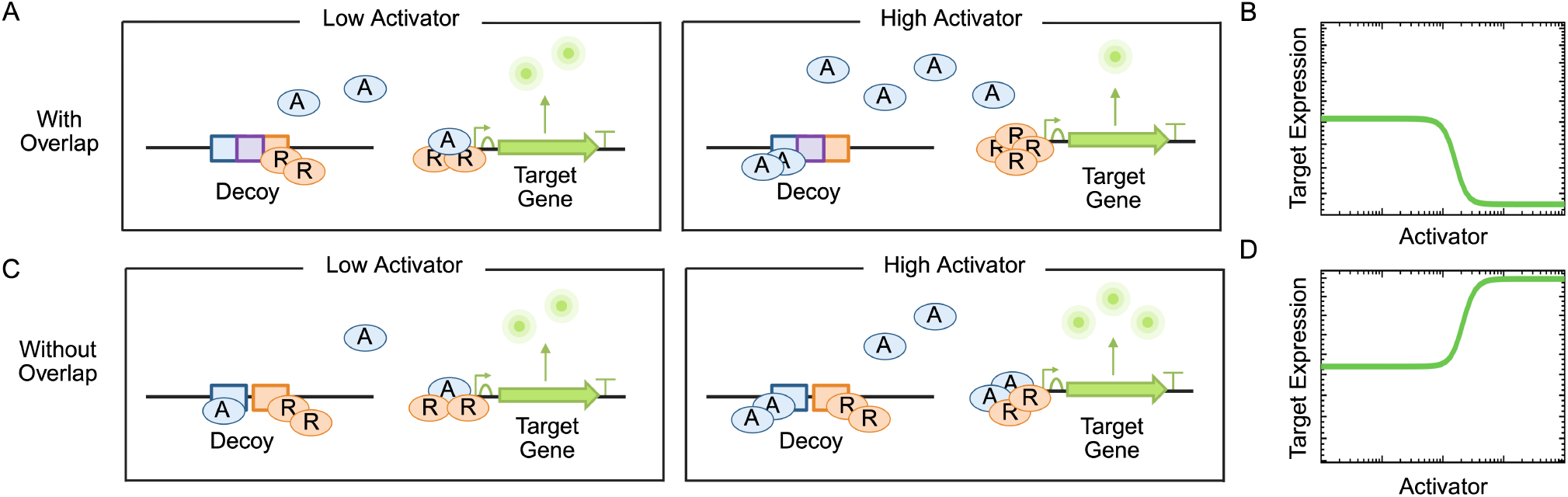
Competition for decoy sites between an activator and a repressor leads to paradoxical target gene regulation. (A) Proposed mechanism for paradoxical gene regulation, where increasing activator level leads to target repression. (B) Paradoxical target repression by the activator predicted by a chemical reaction model considering competitive binding of the regulators at the decoy (SI Appendix, Section 1). (C) When the decoy sites do not overlap, activator and repressor can bind concurrently, and thus increasing activator level does not displace the repressor from the decoy sites. (D) Target expression increases with the activator level as predicted by a chemical reaction model considering independent binding of the regulators at the decoys (SI Appendix, Section 1).

Although implemented in a simplified bacterial context, similar paradoxical behaviors observed in mammalian gene regulation and cancer biology provide a complementary biological context for this study. Metastasis remains the leading cause of cancer mortality, and the epithelial mesenchymal transition plays a central role in enabling tumor cells to acquire migratory and drug resistant phenotypes. The transcription factor *ZEB1* serves as a key EMT regulator and responds sensitively to perturbations in chromatin regulatory networks. Genome wide perturbation studies have revealed that chromatin complexes such as PRC2 and the COMPASS family member KMT2D can exhibit paradoxical influences on EMT gene expression, where removal of an activating complex can lead to higher, not lower, expression of certain targets (16). Recent theoretical work proposed that such counterintuitive responses may arise from competition among epigenetic regulators for shared genomic binding sites, suggesting a resource redistribution mechanism consistent with retroactivity-based models (17). Despite these insights, direct experimental tests remain difficult in mammalian chromatin, where thousands of potential binding sites, dynamic accessibility states, and multilayered feedback loops obscure causal mechanisms. The minimal synthetic system developed here decouples competitive binding from chromatin remodeling and provides a controlled experimental realization of the theoretical competition framework. By demonstrating how simple activator and repressor competition can generate paradoxical regulatory outcomes, this work offers a mechanistic basis for interpreting related behaviors observed during EMT and cancer progression.

## Results

To probe the counterintuitive possibility that higher concentrations of an activator can down-regulate its own target, we analyzed a reaction-network model incorporating competitive binding of an activator and a repressor to decoy binding sites (SI Appendix, Section 1). In the model, the activator (A) and repressor (R) molecules transcriptionally regulate the expression of a target gene while also binding to decoy sites competitively. These decoy binding sites, in turn, sequester the regulators. As the repressor and the activator cannot be bound simultaneously at the decoy sites, increasing the activator level displaces the repressor from the decoy binding sites. This repressor thus is free to bind and repress the target gene (Figure 1A).

In the low-activator regime, the decoy sites are bound mostly by the repressor R. As the concentration of the activator A increases, activator molecules displace the repressor R from the decoys. As a consequence, R becomes available to bind to the target, leading to its repression and the target gene expression decreases as the activator concentration rises (Figure 1B). This paradoxical effect is not observed when the binding to the decoys is independent, that is, when both the activator and the repressor can concurrently bind to the decoys without affecting each other’s ability to bind (Figure 1C). In fact, in this case, increased activator level will only result in more activator bound to the decoys without causing the unbinding of the repressor. More activator also binds to the target and thus increases the target expression (Figure 1D). Thus, competitive binding of A and R to decoy sites is a critical requirement that enables activator-dependent displacement of the repressor and gives rise to paradoxical regulation. With competitive binding, it is therefore also the case that decreasing the activator concentration leads to an increase in target expression due to repressor molecules being sequestered away from the target to the decoy sites.

We built an experimental system in bacterial cells to test the theoretical predictions of Figure 1. We sought a system with an activator and a repressor for which it is possible to independently design decoy binding sites that are overlapping. To this end, we used CRISPR-based gene regulation, with CRISPR activation (CRISPRa) (19, 20) to engineer the activator and CRISPR interference (CRISPRi) (21, 22) to engineer the repressor (Figure 2A). In CRISPRa, an RNA-binding protein fused to an activation domain (RBP-AD) and catalytically inactive Cas9 (dCas9) are both recruited to the DNA by a scaffold RNA (scRNA). Once bound to the DNA, the RBP-AD interacts with the RNA polymerase (RNAP), enhancing the rate of transcriptional initiation (Figure 2A - left) (19). During CRISPRi, dCas9 binds to a guide RNA (gRNA) forming a dCas9/gRNA complex. This complex then binds to the target DNA sequence, physically obstructing RNAP, thereby repressing transcription (Figure 2A - right). By employing CRISPRi and CRISPRa simultaneously, we can achieve independent control of activation and repression of gene expression.

**Fig. 2.**
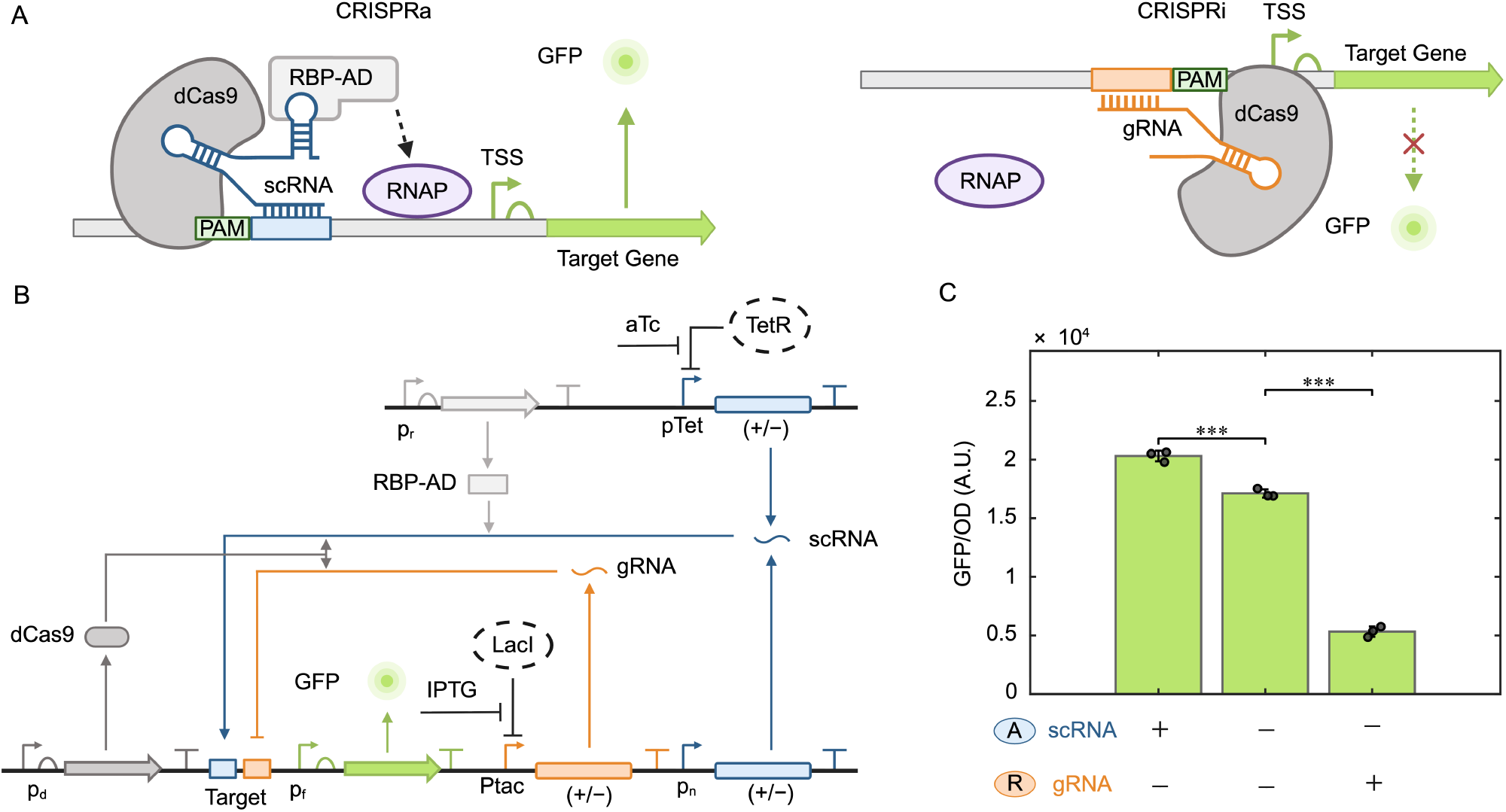
CRISPRa and CRISPRi for combinatorial regulation of a target gene in the absence of decoy sites. (A) (Left) Diagram of the CRISPRa complex consisting of dCas9, RBP-AD, and scRNA binding upstream of the transcription start site (TSS) of the target gene, enhancing gene expression. RBP-AD recruits RNAP, increasing transcription of the target gene. (Right) Diagram of the CRISPRi complex consisting of dCas9 and gRNA binding near the promoter of the target, physically obstructing RNAP binding. Protospacer adjacent motif (PAM) sites are shown for both CRISPRa and CRISPRi complexes. (B) Genetic construct of the target gene (GFP) activated by the scRNA through CRISPRa and repressed by the gRNA through CRISPRi. Expression of gRNA and scRNA is controlled by pTac and Ptet promoters, respectively, with their cognate repressors (LacI and TetR) chromosomally integrated (Marionette strain (18)). (C) Data showing GFP levels for the indicated combinations of the scRNA and gRNA. Here “+ scRNA” refers to 100 nM aTc and the presence of a second scRNA cassette with J119 constitutive promoter (denoted as p_n_), whereas “+ gRNA” corresponds to 10 *µ*M IPTG. “*−* scRNA” corresponds to the absence (including the promoter, gene sequence and terminator) of both scRNA cassettes, while “*−* gRNA” similarly corresponds to the complete removal of the gRNA expression cassette (see SI Appendix, Section 2, Fig. S1, and Tables S2-5 for the plasmid sequences). Error bars represent one standard deviation computed from a set of three independent biological replicates. Every point is computed in arbitrary units (A.U.) (refer to SI Appendix, Fig. S2 for temporal evolution of GFP and a representative OD profile, as OD trajectories were qualitatively similar across all experiments).

To achieve combinatorial regulation of a target gene, we built the genetic construct shown in Figure 2B. Here, the target gene encodes green fluorescent protein (GFP), and the gRNA is expressed by an IPTG-inducible promoter. The scRNA is expressed by both an aTc-inducible promoter and by a constitutive promoter. The scRNA is expressed from two different cassettes to achieve sufficiently high levels of activator. dCas9 and RBP-AD are expressed by a constitutive promoter. Both the activator (scRNA) and the repressor (gRNA) are designed to bind to separate 20bp binding sequences upstream of the transcription start site (TSS) of the GFP expression cassette (Figure 2B). The scRNA target site was placed between positions −81bp and −61bp relative to the TSS, based on prior studies showing that CRISPR-mediated activation is most effective within this window (19). On the other hand, the gRNA target site was positioned between −58bp and −38bp to enable efficient repression while avoiding direct overlay with the target promoter (22). We measured the activated (“+ scRNA” and “*−*gRNA”), basal (“*−*scRNA” and “*−*gRNA”) and repressed (“*−*scRNA” and “+ gRNA”) expression of GFP. Here, “+ scRNA” represents the presence of both scRNA expression cassettes with maximum aTc induction, while in “*−*scRNA” both scRNA-expressing cassettes are removed. As expected, the GFP levels are higher than basal when activated, whereas they are lower than basal when repressed (Figure 2C, corresponding model data is shown in SI Appendix, Fig. S3). We then designed the decoys such that the binding sites for the scRNA and gRNA overlap, thereby leading to competitive binding of the activator and repressor. The distance of the decoy sites from the TSS is chosen with consideration to the optimal binding site positions for CRISPRi and CRISPRa found in (22) and (19), respectively. In particular, we placed the scRNA binding site between the −81bp and the −61bp sites with respect to the TSS for optimal activation (19). The gRNA binding site is placed in the −71bp to −51bp region to intentionally overlap with the scRNA binding region by 10bp. The PAM sites (indicated as AGG) for each of the CRISPRi and CRISPRa complexes flank either side of this 30bp region (Figure 3A). To verify that the activator displaces the repressor from the decoys in order to bind, we employed a genetic construct where the decoy sites are placed upstream of a promoter expressing a red fluorescent protein (RFP) (Figure 3B). We then first assessed the ability of the scRNA to activate RFP expression and of the gRNA to repress RFP expression. Specifically, we verified repression by the gRNA by achieving a lower RFP expression level in the presence of the gRNA and absence of the scRNA (“+ gRNA” and “−scRNA”) compared to the basal RFP level obtained in the absence of both scRNA and gRNA (“−scRNA” and “-gRNA”) (Figure 3C). Similarly, we verified activation by the scRNA by achieving a higher RFP expression in the presence of the scRNA and absence of the gRNA (“+ scRNA” and “-gRNA”) compared to the basal RFP level (Figure 3C).

**Fig. 3.**
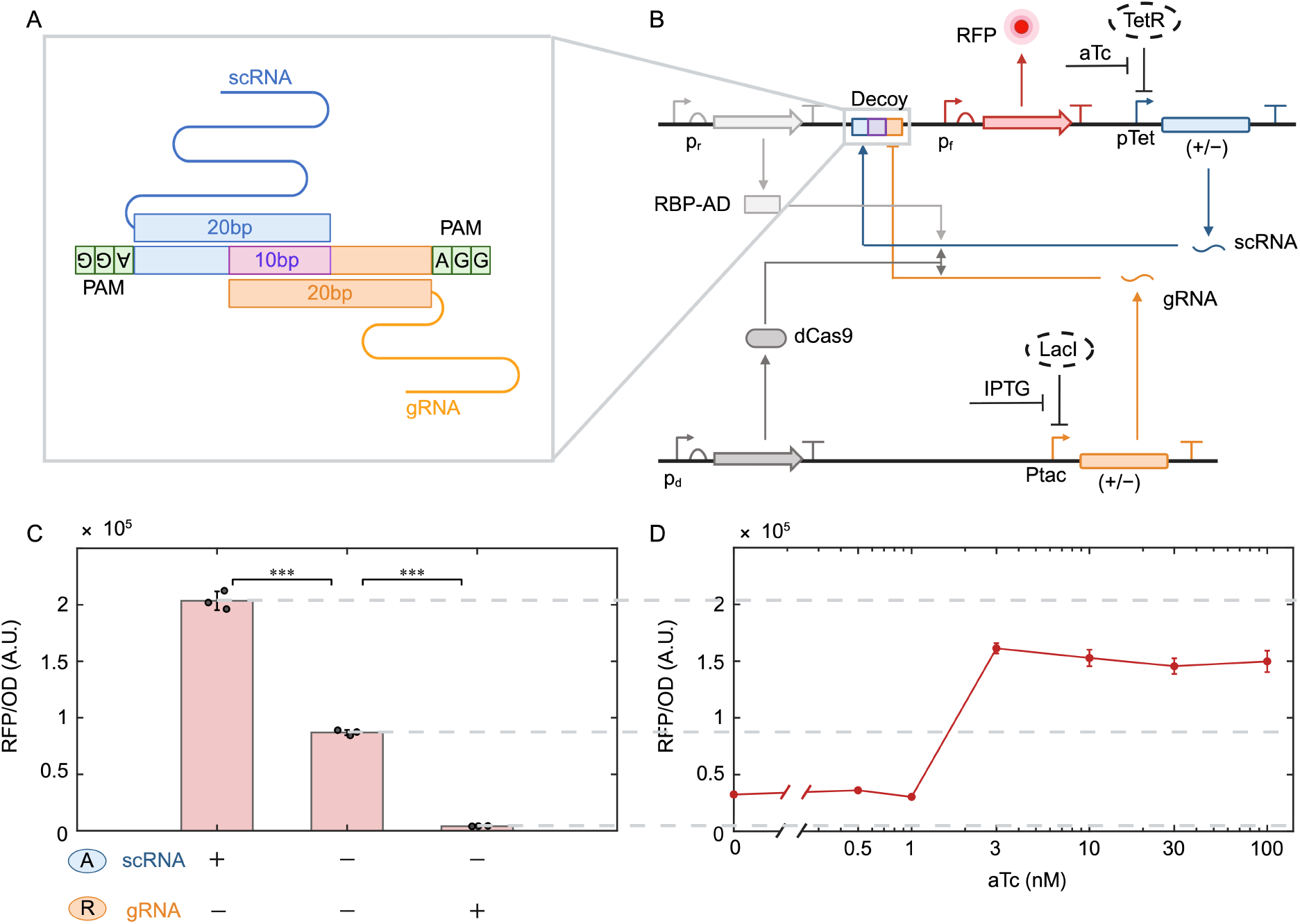
Competitive binding at decoy sites is implemented via overlapping binding sites for scRNA and gRNA. (A) Genetic diagram showing the implementation of overlapping binding sites between the scRNA and the gRNA at the decoy (see SI Appendix, Section 2, Fig. S1, and Tables S2-5 for the plasmid sequences). (B) Genetic construct with RFP placed downstream of the decoy sites activated by the scRNA and repressed by the gRNA. (C) Data showing RFP levels for varying combinations of the scRNA and gRNA. Here “+ scRNA” refers to 100 nM aTc, “+ gRNA” corresponds to 10 *µ*M IPTG and “*−* scRNA” or “*−* gRNA” corresponds to the lack of scRNA or gRNA gene sequence. (D) Response of RFP to increasing scRNA (through aTc induction) in the presence of repressor (10 *µ*M IPTG). The dashed lines are the extension of the respective values from panel (C). All plasmids are introduced into the Marionette *E. coli* strain (18), where the TetR and LacI regulators are endogenously expressed. Error bars represent one standard deviation computed from a set of three independent biological replicates. Every point is computed in arbitrary units (A.U.) (refer to SI Appendix, Fig. S4 and S5 for RFP time series).

To verify whether the activator would displace the repressor from the decoys, we varied the activator scRNA level in the presence of the repressor gRNA. We thus set the gRNA level to a constant value by a fixed level of IPTG such that the RFP expression was lower than the basal value (indicated by the central dashed line in Figure 3D) and increased the scRNA level through aTc concentration. As the amount of scRNA is progressively increased, RFP level increases crossing the basal expression level, but remaining below the maximally activated level (indicated by the upper dashed line in Figure 3D). The ability of the scRNA to recover RFP expression above the basal level suggests that the scRNA competes with the gRNA for binding at the decoy sites, as expected from competitive binding. Specifically, the increase in RFP expression despite the continued presence of the repressor gRNA suggests that increasing scRNA levels displace repressors from the decoy sites. If the repressors remained bound at the same occupancy, RNAP obstruction would persist and RFP expression would remain repressed below the basal level. The corresponding model predictions are shown in SI Appendix, Fig. S3.

To assess whether paradoxical target regulation could be observed, we combined the target with the designed decoy sites into a new genetic construct (Figure 4A). This construct allows constant expression of the repressor, tunable levels of the activator, and overlapping binding sites at the decoy by combining the constructs in Figure 2B and Figure 3B. We observed that as the activator level is increased by increasing the scRNA level via aTc, GFP levels decrease by approximately 40% showing an unintended target repression (Figure 4B). The same experiment performed using a functional RBS upstream of the RFP decoy reporter is shown in SI Appendix, Fig. S6, where both paradoxical repression of GFP and activation of the RFP decoy reporter are observed, consistent with activator binding at the decoy sites (SI Appendix, Fig. S6).

**Fig. 4.**
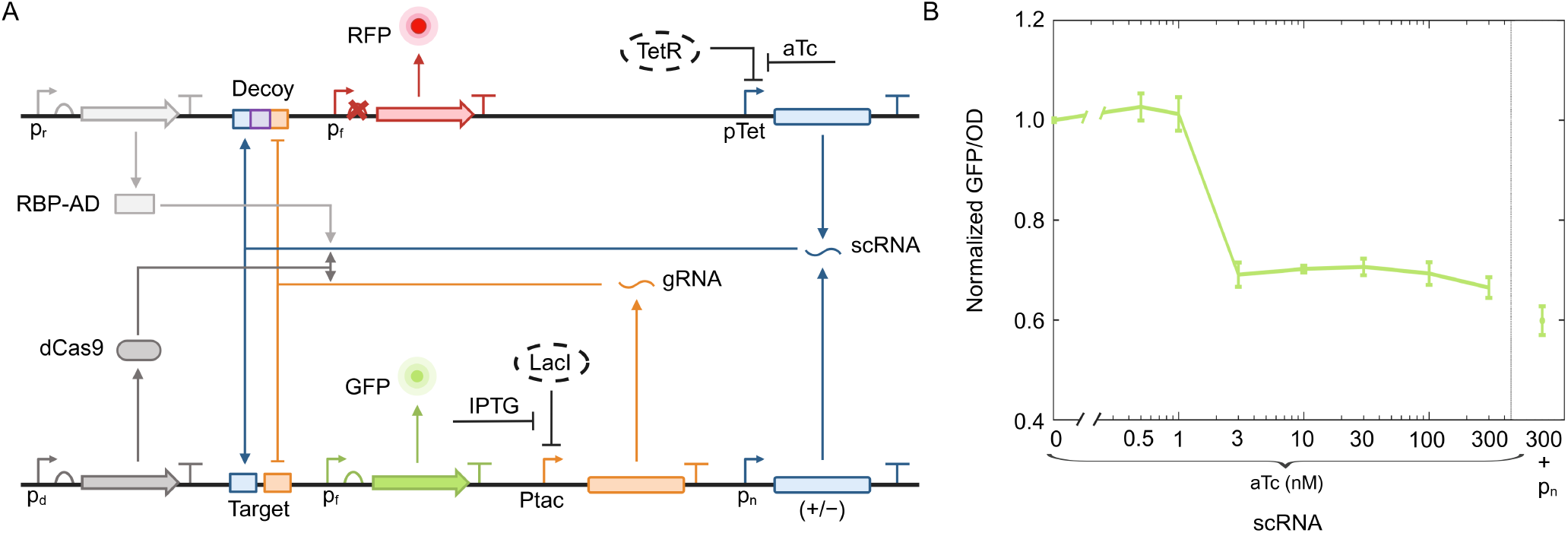
Paradoxical repression of the target gene by the activator. (A) Genetic construct built where the target and decoy binding sequences are the same as in Figure 2 and 3. The level of gRNA is kept constant with 10 *µ*M IPTG. The scRNA is produced from two distinct cassettes, one with an aTc inducible promoter and the other with a constitutive J119 promoter. The TetR and LacI regulators are expressed from the chromosome (Marionette strain (18)). The ribosome binding site (RBS) of the RFP cassette downstream of the decoy sites is changed to a dead RBS (see SI Appendix, Table S4 for sequences and Fig. S6 for data with functional RBS for RFP) to prevent competition for translational resources (23). (B) Output of target gene (normalized GFP) with increasing scRNA. The GFP values are normalized using the GFP level at 0 nM aTc (i.e. no induction) and without the secondary scRNA expression cassette. The scRNA level is increased through aTc concentration as indicated, with or without the secondary scRNA expression cassette (J119) as indicated. Error bars represent one standard deviation computed from a set of three independent biological replicates (refer to SI Appendix, Fig. S7 for GFP temporal data).

Next, we investigated the effect of the binding affinity of the activator to the target. The model predicts that the paradoxical effect is observed only when the dissociation constant between the CRISPRa complex and the target binding site (denoted as *K*_*ta*_) is sufficiently large, i.e., when the activator–target binding affinity is weak (Figure 5A). Specifically, for low values of *K*_*ta*_, the target gene is activated by increasing activator levels (Figure 5A-1,2), whereas we observe the paradoxical repression by the activator for high values of *K*_*ta*_ (Figure 5A-3). To experimentally test whether decreased *K*_*ta*_ (increased activator binding affinity to the target) would remove the paradoxical repression of the target by the activator, we decreased the number of mismatches between the scRNA and the DNA binding sequence on the target (Figure 5B). Specifically, starting from the scRNA binding sequence used in Figure 4, which contained three mismatches, we progressively removed mismatches up to 0 mismatches (Figure 5B). With three mismatches, the activation level is low (approximately 1.3 fold), which increases to approximately 4-fold and 5-fold activation as the number of mismatches is decreased to 1 mismatch and 0 mismatches, respectively (Figure 5B). These results confirm the model predictions (SI Appendix, Fig. S3). For the extreme case of complete mismatch between the scRNA and target binding site (20 MM), paradoxical repression is again observed (SI Appendix, Fig. S12).

**Fig. 5.**
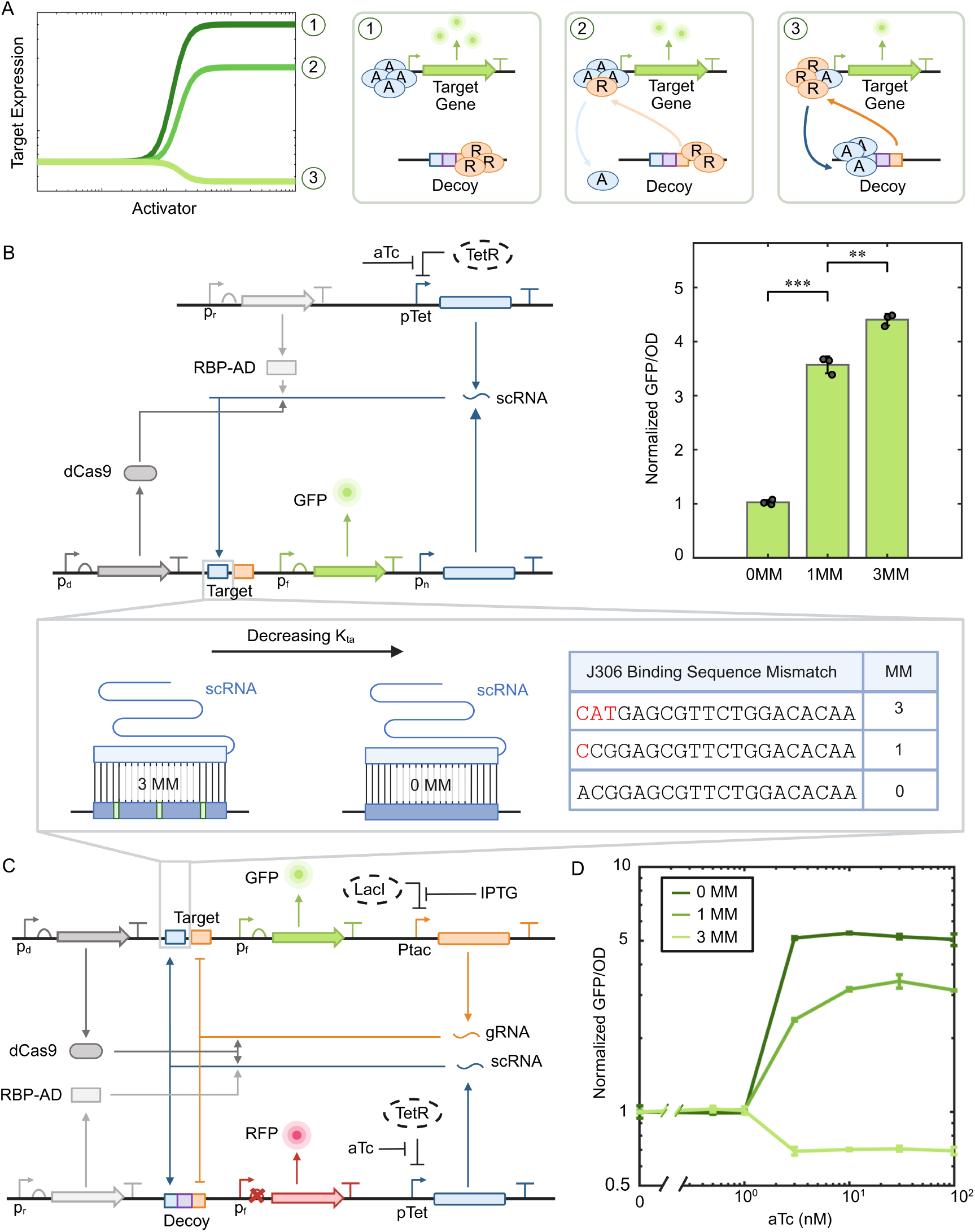
Paradoxical repression is observed for low binding affinity of the activator to the target. (A) Numerical simulations showing the target expression with increased activator level for different values of the dissociation constant of the scRNA to the target (*K*_*ta*_). We observe activation for low *K*_*ta*_ values and paradoxical repression for high *K*_*ta*_ values. (B-Left) Genetic construct with varying *K*_*ta*_ through the variation of mismatches at the activator binding site on the target. (B-Right) Fold activation of GFP for different numbers of mismatches in the target-activator binding site. The fold activation is the ratio of GFP in the presence of both scRNA cassettes with respect to the GFP in the absence of both scRNA cassettes. (C) Genetic construct to investigate the effect of target-activator binding affinity (through removal of mismatches) on the paradoxical repression. (D) Response of normalized GFP to increasing scRNA, for different numbers of mismatches. The GFP values are normalized using the corresponding GFP levels at 0 nM aTc (i.e. no induction). The amount of gRNA is kept constant with 10 *µ*M IPTG and the scRNA levels are varied through aTc as indicated. In (B) and (C), all plasmids are introduced into the Marionette *E. coli* strain (18), where the corresponding regulators are endogenously expressed. Error bars represent one standard deviation computed from a set of three independent biological replicates (refer to SI Appendix, Fig. S8-11 for GFP time series).

Next, we investigated the additional requirements for the paradoxical repression by the activator: the presence of the decoy, competitive binding at the decoy, and the presence of the repressor. To demonstrate the necessity of these elements, we removed them systematically one at a time and observed changes in the regulation of target expression by the activator. In each of the three ablation studies, we started from the configuration in Figure 5C with 3 MM mismatch at the target site. First, we removed only the decoy sites from the genetic construct (Figure 6A). In this configuration, increasing the level of the scRNA results in increasing GFP expression (Figure 6B), thereby indicating a lack of paradoxical regulation. Next, to verify the necessity of the competitive binding at the decoy sites, we modified the decoy sites to be non-overlapping. In particular, the 20bp binding sequence for the repressor was moved downstream of the scRNA cassette such that the activator and repressor binding sites are more than 1000 base pairs apart (Figure 6C). In this system, the repressor cannot be displaced by the activator. The expression of GFP again increases with the scRNA level, showing no paradoxical effect (Figure 6D). The increased activator concentration therefore leads to expected activation, albeit lower than that observed in the no-decoy case due to sequestration of activators by the decoys (compare activation in Figure 6B with no decoy sites and Figure 6D with non-overlapping decoy sites). Finally, we investigated the requirement for the presence of the repressor by removing the gRNA cassette from the genetic construct (Figure 6E). The dose-response curve shows that GFP levels remain relatively constant as the aTc concentration is increased (Figure 6F) showing again that the paradoxical effect was removed. This experiment demonstrates that the paradoxical repression requires the presence of the repressor molecule that is being displaced, consistent with former studies on retroactivity (12). As an additional control, a non-targeting gRNA was used in place of the repressor gRNA, which similarly removed the paradoxical repression (SI Appendix, Fig. S13). Consistent with these findings, the mathematical model predicts the absence of paradoxical repression upon systematic removal of each required genetic component (SI Appendix, Fig. S14).

**Fig. 6.**
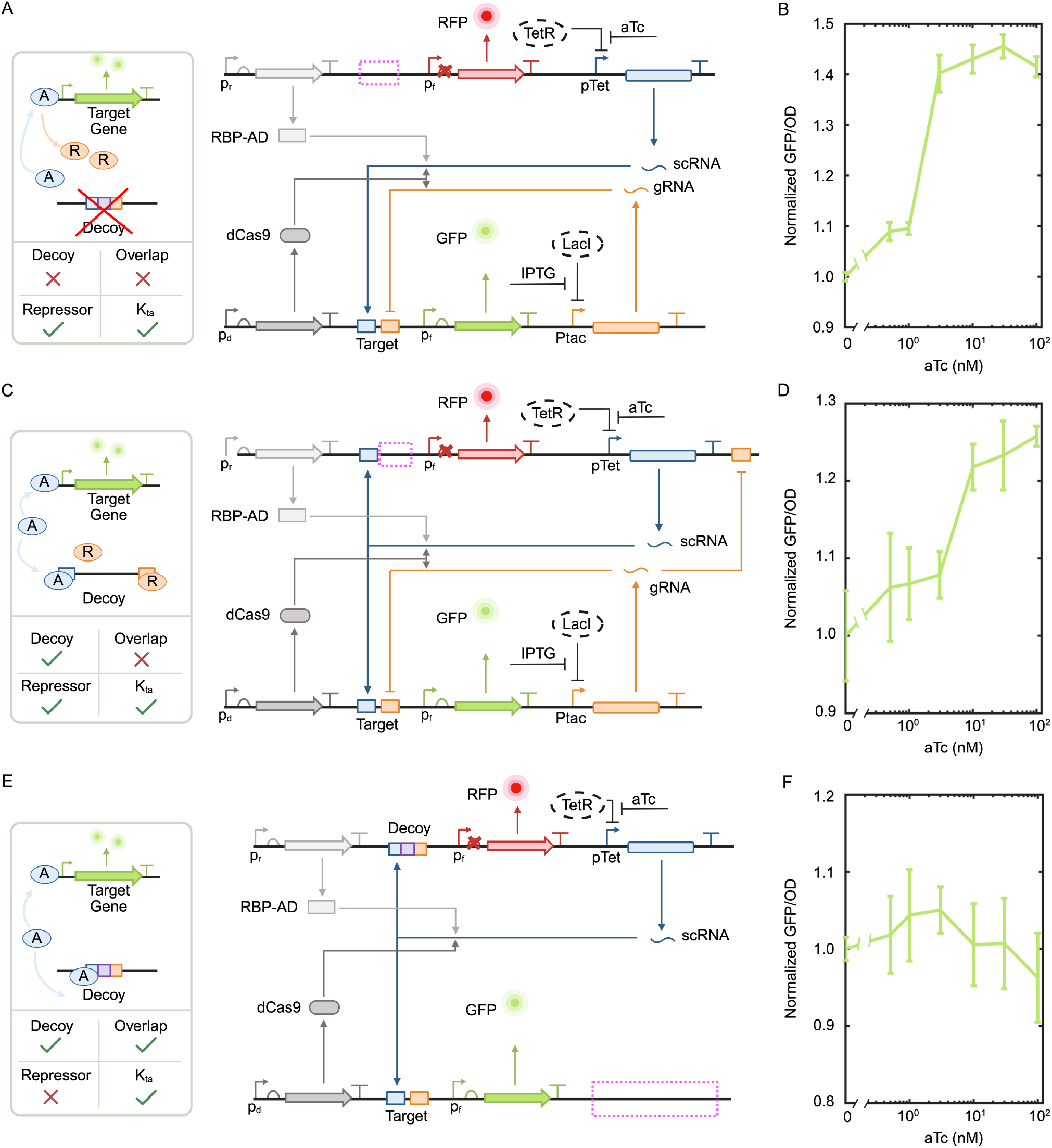
The repressor and overlapping decoy sites are critical to observe the paradoxical behavior. (A) Genetic construct previously used (in Fig. 4) to observe the paradoxical effect with decoy sites removed and with 3 MM mismatch at the target site. (B) Input/output response of GFP with increasing aTc for the construct in (A) for 10 *µ*M IPTG. (C) Genetic construct previously used to observe the paradoxical effect with competitive binding at decoy sites removed by undoing the overlap. (D) Input/output response of GFP with increasing aTc for the construct in (C) for 10 *µ*M IPTG. (E) Genetic construct previously used to observe the paradoxical effect with repressor removed. Input/output response of GFP with increasing aTc for the construct in (E). In (A), (C) and (E), all plasmids are introduced into the Marionette *E. coli* strain (18), where the corresponding regulators are endogenously expressed. In (B), (D) and (F), the GFP values are normalized using the corresponding GFP levels at 0 nM aTc (i.e. no induction). Error bars represent one standard deviation computed from a set of three independent biological replicates. Every point is computed in arbitrary units (A.U.) (refer to SI Appendix, Fig. S15-17 for temporal GFP data and SI Appendix, Fig. S18 for corresponding data with 0, 1, 3 and 20 mismatches at the scRNA target binding site).

## Discussion and Conclusions

Many gene regulatory systems involve multiple effectors acting simultaneously on the same target, yet most mechanistic studies have analyzed regulators in isolation. The present work was motivated by a central question that arises when both an activator and a repressor operate within a shared regulatory environment: how does competition between distinct transcription factors reshape transcriptional responses? Rather than considering a single regulator acting alone, our study focuses on the interplay between positive and negative effectors whose activities are coupled through shared binding sites and limited molecular resources, creating a setting in which redistribution of regulatory activity becomes a dominant determinant of gene-expression outcomes.

Our theoretical and experimental results demonstrate that competition between an activator and a repressor can produce counterintuitive outcomes in which increasing the abundance of a nominally positive regulator leads to reduced expression of the target gene. This effect does not reflect a change in the intrinsic biochemical role of the activator or repressor. Instead, it arises from redistribution of regulators across a common pool of decoy sites, a form of retroactivity in which downstream binding interactions feed back onto regulator availability and reshape effective input signals (12, 14, 24). The mechanistic reaction-network model developed here predicted that activator induction could release repressors from decoy sites and thereby enhance repression at the promoter, a prediction that was validated experimentally using a minimal CRISPRa/CRISPRi circuit. Competitive binding at decoy sites was implemented through overlapping binding sequences, and systematic perturbations confirmed that exclusive binding and target affinity are essential determinants of the paradoxical regime.

Although the primary focus of this work is on dual-regulator competition, these findings can be understood within the broader framework of regulatory duality discussed in the Introduction. Intrinsic models describe how a single transcription factor may exhibit both activating and repressing influences through local TF–RNAP interactions, whereas extrinsic models emphasize how resource competition reshapes effective regulatory inputs (2, 6, 7). The present results extend these ideas by showing that paradoxical responses can emerge purely from interactions between distinct regulators, even when their intrinsic biochemical roles remain unchanged. In this sense, multi-factor competition provides a minimal mechanistic route to behaviors often attributed to more complex network-level effects and highlights how coupling between regulators can modify the effective gain of transcriptional control.

More broadly, the synthetic platform developed here illustrates how binding competition redistributes functional states across coupled molecular components and thereby modulates the effective gain of gene regulation. By decoupling competitive binding from chromatin remodeling and higher-order feedback, the system isolates a core principle that may operate across diverse biological contexts. The results highlight how nonlinear competition can amplify or invert apparent regulatory outcomes without altering the underlying sign of individual molecular interactions, reinforcing the view that network context plays a central role in shaping transcriptional behavior.

Paradoxical regulatory responses of this kind are not unique to transcriptional networks. Similar activation-by-inhibition effects have been observed in cooperative enzymatic systems, where partial occupancy by an inhibitor transiently enhances catalytic output before suppressing it at higher concentrations (25). In such systems, partial inhibitor binding shifts the balance between inactive and active conformations and effectively enhances the propensity of unoccupied sites to engage in productive ligand interactions. Conceptually, this mechanism parallels duality in gene regulation: just as a transcription factor can simultaneously stabilize RNAP binding while slowing transcriptional initiation, or redistribute repressors through competition for decoy sites, partial binding in an enzyme complex can improve the catalytic activity of the remaining unbound sites. These parallels suggest that activation-by-repression and activation-by-inhibition may reflect a shared phenomenology of nonlinear biochemical systems in which binding events reshape system dynamics rather than acting as purely monotonic inputs. Such paradoxical effects have also been documented in kinase signaling networks. In the mitogen-activated protein kinase pathway, clinically used inhibitors targeting wild-type BRAF can promote dimerization with unliganded CRAF, relieving autoinhibition and enhancing downstream signaling activity (26, 27). Although the molecular details differ from transcriptional regulation, the underlying logic is similar: an interaction that appears inhibitory at the local level can increase system-level output by redistributing activity across coupled components. Together with the enzymatic example, these observations point toward a unifying principle across biological scales, in which competition and allosteric coupling generate nonlinear input–output relationships.

While the synthetic circuit studied here does not reproduce the full complexity of eukaryotic chromatin regulation, it provides a controlled framework for testing mechanisms proposed to explain paradoxical responses in cancer epigenetics (17). Metastasis accounts for most cancer-associated mortality (28), and epithelial-to-mesenchymal transition (EMT) is a key process (29). Although EMT is often framed in terms of transcription factors, epigenetic regulators are key determinants of EMT stability and plasticity (30, 31). One key transcription factor, *ZEB1*, is especially sensitive to upstream epigenetic perturbations that affect it or its regulators, such as miR-200 (32–36).

Antagonism of chromatin modulators is known to regulate the EMT (37). As an example, PRC2-dependent H3K27 methylation helps restrain mesenchymal programs and support epithelial-state residence in multiple carcinoma contexts (38– 40). COMPASS (Complex of Proteins Associated with Set1) is a central player in epigenetic regulation, but COMPASS-related regulation is context dependent: depending on lineage state and network configuration, these perturbations can either constrain or facilitate EMT-associated programs (41, 42). This context dependence argues against assigning fixed state-protective roles to single complexes across tumor types. For instance, genome-wide perturbation studies further show that PRC2 and KMT2D-COMPASS loss does not produce a uniform monotonic shift in expression. For *ZEB*1, promoter occupancy and expression responses can change non-intuitively under perturbation; in some settings, loss of an activating factor (KMT2D) is accompanied by increased *ZEB1* expression (41). Other studies have supported context-dependence in setting of regulator antagonism in a variety of epigenetic systems (43–46). These observations highlight a broader interpretation challenge for chromatin-perturbation experiments: regulators act on overlapping genomic territories and can redistribute across shared target pools. Although the synthetic system does not capture nucleosome dynamics or histone modification feedback, it provides a simplified testbed to generate hypotheses and study how redistribution of regulators across shared binding pools can generate paradoxical expression changes similar to those observed during EMT and tumor progression.

In summary, this study demonstrates that competition between an activator and a repressor for overlapping binding sites can generate paradoxical gene-expression responses through retroactivity-mediated redistribution of regulatory activity. By linking intrinsic regulatory mechanisms with extrinsic resource competition, and by connecting synthetic circuit experiments to broader biochemical and signaling examples, our results suggest that nonlinear redistribution of molecular activity may represent a general organizing principle of biological regulation. Viewed through this lens, activator– repressor competition provides not only a minimal explanation for unexpected transcriptional behaviors but also a conceptual bridge connecting gene regulation, enzymatic control, and signaling dynamics across diverse biological systems.

## Methods

### Bacterial strain and growth medium

Plasmids used in this work were assembled using standard molecular cloning techniques. Genetic constructs were transformed in the bacterial strain *E. coli* NEB5*α* (NEB, C2987H) grown in LB Broth Lennox (SIGMA-ALDRICH, L3022-1KG) supplemented with appropriate antibiotics. Following sequence verification, plasmids are co-transformed to *E. coli* Marionette strain (18) for experiments presented in this study. Experiments were performed in M9 minimal medium containing M9 salts (1X), 0.4% glucose (SIGMA-ALDRICH, 49159), 0.2% casamino acid (VWR, TS61204-5000), 1 mM thiamine hydrochloride (SIGMA-ALDRICH, T4625-100G), magnesium sulfate (NEB, B1003S), and calcium chloride (SIGMA-ALDRICH, C4901-100G) with appropriate antibiotics. Ampicillin (SIGMA-ALDRICH, A0166-5G) and kanamycin (SIGMA-ALDRICH, K1377-5G) were used at final concentrations of 100, and 50 *µ*g mL^*−*1^, respectively. Isopropyl *β*-D-1-thiogalactopyranoside (IPTG, SIGMA-ALDRICH, I6758-5G) and anhydrotetracycline (aTc, SIGMA-ALDRICH, 37919-100MG-R) are used as the inducers.

### Cloning

The genetic cloning was based on Gibson assembly using plasmid 1 and plasmid 2a from Aravind et al. (20) as backbone vectors. DNA oligos were synthesized using the Oligo Rapid service provided by Azenta Life Sciences. DNA fragments for assembly were amplified by PCR using Q5 High-Fidelity 2X Master Mix (NEB, M0492L). PCR products underwent gel electrophoresis for purification and the gel was extracted using the Zymoclean Gel DNA Recovery Kit (Zymo Research, D4002). Purified DNA fragments were subsequently assembled using the Gibson Assembly Master Mix (NEB, E2611L). The assembled plasmids were transformed into chemically competent cells, and single colonies were inoculated into LB medium and cultured overnight at 37°C. Plasmid DNA was extracted from overnight cultures using the ZR plasmid miniprep-classic kit (Zymo Research, D4015). DNA sequencing was performed by Primordium Labs. The plasmids and genetic parts are listed in SI Appendix, Section 2, Fig. S1 and Tables S2-5.

### Plate reader experiments

Frozen glycerol stocks stored at −80°C were used to inoculate overnight cultures. Cultures were grown in M9 medium supplemented with appropriate antibiotics at 37 °C, shaking at 250 rpm using a horizontal orbiting shaker for 13-16 hours in 15 mL culture tubes (VWR, 60818-667). These overnight cultures were diluted to an initial optical density (OD) of 0.01 in 200 *µ*L of growth medium per well in Falcon 96-well cell culture plates with a flat bottom (VWR, 15705-066) containing appropriate antibiotics and inducers. Optical density was calculated from the measured absorbance at 600 nm (*OD*_600_) after subtracting the background contribution (*OD*_*M* 9_) from M9 minimal medium (*OD* = *OD*_600_*− OD*_*M* 9_). Plates were incubated at 30°C in a Tecan Infinite 200 PRO microplate reader under static conditions. Prior to each OD600 and fluorescence measurement, plates were shaken for 5 s at ‘fast’ speed. Measurements were acquired at 5 min intervals. RFP fluorescence was monitored using excitation and emission wavelengths of 584 and 619 nm, respectively, with a gain of 100 while GFP fluorescence was measured using excitation and emission wavelengths of 485 and 530 nm, respectively, with a gain of 70.

To maintain exponential growth, cultures were diluted with fresh growth medium supplemented with antibiotics and inducers to an OD of 0.01 when the OD approached 0.08 at the end of each batch. Multiple serial dilution batches were continued until gene expression reached steady state. Reported steady-state fluorescence values were recorded when cultures reached an OD of 0.06 during the final batch.

### Data Analysis

Raw measurements of GFP fluorescence (*GF P*_*raw*_), RFP fluorescence (*RFP*_*raw*_), and optical density at 600 nm (*OD*_600_) were obtained directly from the plate reader. The optical density of each sample was background-corrected by subtracting the optical density contribution of the M9 minimal medium (approximately 0.08):

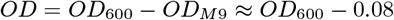

To normalize fluorescence signals by cell density, GFP and RFP values were first corrected by subtracting their corresponding background fluorescence values and subsequently divided by the background-corrected OD value:

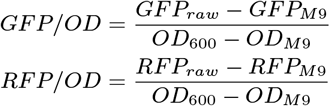

where *GF P*_*M* 9_ and *RFP*_*M* 9_ denote the corresponding background fluorescence values of the M9 medium.

For the data shown in Figures 4, 5, and 6, normalization of GFP was performed with respect to the corresponding GFP levels at 0 aTc (no induction). To normalize a point (*m, s*) (where *m* is the mean and *s* is the standard deviation), we computed the new point 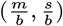 where *b* is the corresponding mean GFP level at 0 aTc (no induction).

### AI use

During the preparation of this manuscript, the authors used ChatGPT (GPT-5, OpenAI) to improve the clarity and readability of the Introduction and Discussion sections.

## Acknowledgments

This work was partially supported by grants AFOSR FA9550-22-1-0316 and NSF CCF FET Award 2007674

## Supporting Information Text

### 1. Chemical reaction model of the system

To guide our prototyping, design relevant experiments, and isolate the primary drivers of the paradoxical regime, we developed a streamlined mathematical model rather than an exhaustive analytical one. For this purpose, we started from a chemical reaction model of transcriptional regulators for genomic binding sites developed previously in (1), using a set of parameters that fit our experimental measurements.

Here, we review the chemical reaction model used in (1). We consider a target gene (*T*) that produces the output protein (*Y*) at a basal production rate, *β*. When an activator (*A*) binds to the target, the gene enters an active state (*T*_*A*_), and produces the protein at an increased rate, *κ > β*. The target can also bind to a repressor (*R*) placing the gene in a repressed state (*T*_*R*_).

The reactions involved are as follows:

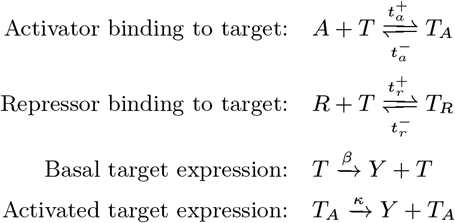

The activator and repressor can also bind to other genomic sites, where they compete for binding. We model these additional sites as decoys (*D*). A decoy bound to an activator is denoted by *D*_*A*_, while a decoy bound to a repressor is denoted by *D*_*R*_.

The reactions involved are as follows:

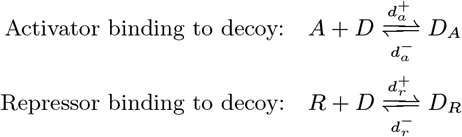

The remaining reactions describe the production of the activator and repressor, as well as degradation. Here, *u*_*A*_ and *u*_*r*_ denote the production rates of the activator and repressor, respectively, and *δ* denotes the degradation rate. The reactions are as follows:

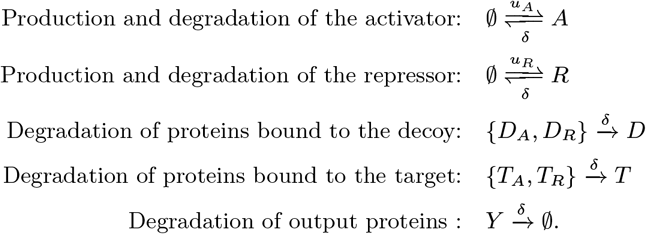

Using mass-action kinetics, the above reactions can be expressed as a system of reaction-rate equations used to simulate the dynamics of the system:

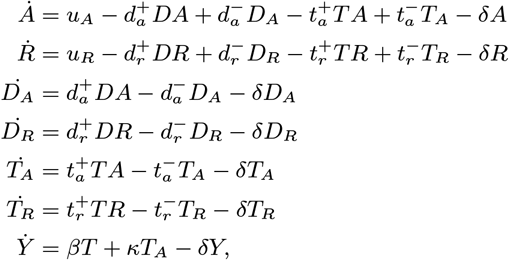

We define the total concentrations of the activator (*A*_*t*_), repressor (*R*_*t*_), target (*T*_*t*_), and decoy (*D*_*t*_) species as:

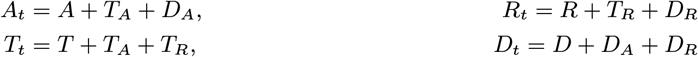

#### A. Steady-state behavior of the system

At steady state, all time derivatives are set to zero, yielding:

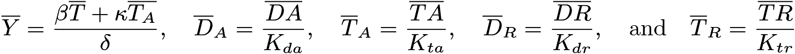

Where 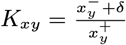. Substituting in the total concentration equations, we get:

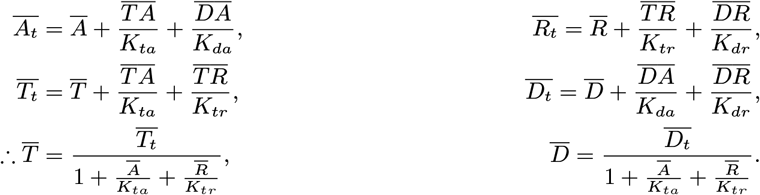

Combining the above equations, we get:

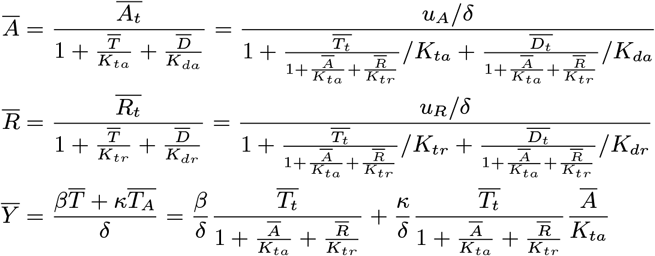

The production of the activator using the aTc-inducible pPhlF promoter is modeled as:

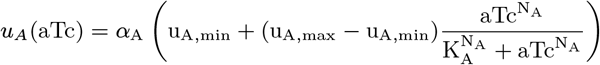

and the production of the repressor using the IPTG-inducible Ptac promoter is modeled as:

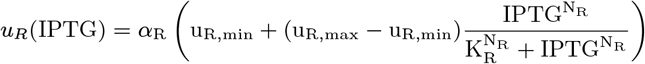

#### B. Parameter estimation

To estimate the model parameters, we used MATLAB’s fmincon function to find one set of parameters that approximately minimizes an objective function, defined as the sum of the squared differences between the experimental data and the fitted model outputs under equivalent experimental conditions. The following experimental data sets were used for the fitting process:

- GFP data from Figure 2(C)
- RFP data from Figure 3(C-D)
- GFP data from Figure 5(B)

The parameters fitted from the model are shown in Table S1.

Other parameter sets may fit the data equally well, but the qualitative conclusions should remain unchanged. We intentionally used a minimal mathematical model to capture the core mechanics of the paradoxical regime. Our goal was not a high-fidelity quantitative simulation, but rather a practical framework to guide system construction and predict key experimental parameters. Nonetheless, the model was useful in predicting key experimental behaviors. Parameter fitting was performed using only a limited subset of experimental data: target activation, basal, and repression levels in the absence of decoy sites (Figure 2C); decoy activation, basal, and repression levels in the absence of target sites (Figures 3C-D); and target activation across different mismatch levels in the absence of repressors or decoy sites (Figure 5B). The numerical simulations corresponding to the data used for parameter fitting are shown in Figure S3.

To test the predictive capability of the model, we then generated simulations for the paradoxical gene regulation. For example, comparison of Figures 5(A) and 5(D) demonstrates that the model predictions were qualitatively validated. We further used the model to predict that paradoxical repression would disappear upon removal of the key requirements (simulated loss of behavior shown in Figure S14). Experimental validation confirmed this prediction, as the paradoxical behavior indeed disappeared when the requirements were removed individually (Figure 6).

Note: parameters such as total DNA (copy number), degradation rate, and the ones for aTc induction are obtained from the appropriate references.

**Table S1.** Parameters obtained through fitting.

| Parameter | Symbol | Parameter Values | Reference |
| --- | --- | --- | --- |
| Total concentrations: | $T_t$ (Target gene) | 15 nM for P15A | (2, 3) |
| | $D_t$ (Decoy sites) | 80 nM for PSC101(E93G) | (2, 3) |
| Activated Production Rate | $\kappa$ | 240 $\text{min}^{-1}$ | experiment |
| Basal Production Rate | $\beta_1$ | 30 $\text{min}^{-1}$ | experiment |
| Degradation rate | $\delta$ | $\frac{\log(2)}{20} \text{min}^{-1}$ | (4) |
| aTc Induction: | $\alpha_A$ | 250 $\text{min}^{-1}$ | experiment |
| | $K_A$ | 2 nM | experiment |
| | $N_A$ | 4 | (5) |
| | $u_{A,min}$ | 0.0005 nM | (5) |
| | $u_{A,max}$ | 2.5 nM | (5) |
| IPTG Induction: | $\alpha_R$ | 40 $\text{min}^{-1}$ | experiment |
| | $K_R$ | 14 nM | experiment |
| | $N_R$ | 2 | (5) |
| | $u_{R,min}$ | 0.005 nM | (5) |
| | $u_{R,max}$ | 3.3 nM | (5) |
| Dissociation constants: | $K_{tr}$ | 15 nM | - |
| | $K_{dr}$ | 15 nM | - |
| | $K_{da}$ | 50 nM | - |
| | $K_{ta}$ (for 0MM) | 50 nM | - |
| | $K_{ta}$ (for 1MM) | 200 nM | - |
| | $K_{ta}$ (for 3MM) | 12000 nM | - |

### 2. Plasmid maps and Sequences

All genetic constructs used in this study were derived from two parent plasmids, hereafter referred to as Plasmid 1 and Plasmid 2. Unless otherwise noted, all constructs described in the manuscript were generated through minor sequence modifications to these two backbone designs, while the remaining plasmid architecture was kept unchanged.

Plasmid 1 is a p15A-based vector (approximately 15 copies per cell) carrying kanamycin resistance (Figure S1A). This plasmid constitutively expresses dCas9 and contains a gRNA expressed from an inducible Ptac promoter. The target reporter gene, GFP, is positioned downstream of the scRNA and gRNA binding region. To investigate the effects of target activator binding, the scRNA binding site located upstream of GFP was modified to contain varying numbers of mismatches relative to the corresponding guide sequence. Specifically, constructs containing 0 mismatch (0 MM), 1 mismatch (1 MM), and 3 mismatches (3 MM) were generated. These binding-site sequences are shown in Figure 5(B) of the main manuscript. In the constructs used for Figures 2, 4, and 5(B) of the main manuscript, an additional constitutively expressed scRNA cassette was inserted downstream of the gRNA cassette. Aside from these specified modifications, the remainder of Plasmid 1 was identical across all constructs used in the study.

Plasmid 2 is based on the pSC101(E93G) origin (approximately 80 copies per cell) and carries ampicillin resistance (Figure S1B). This plasmid constitutively expresses the RBP–AD fusion protein and contains an aTc-inducible scRNA cassette. In addition, overlapping decoy binding sites were positioned upstream of an mRFP reporter containing a nonfunctional (“dead”) RBS. The overall architecture of Plasmid 2 remained unchanged for all experiments shown in Figures 1–5.

For the constructs used in Figure 6(A), the overlapping decoy sites were removed by deleting the J306 and gRNA binding sites located upstream of the mRFP reporter. For Figure 6(C), non-overlapping decoy sites were generated by relocating the gRNA binding site originally positioned upstream of mRFP to a location downstream of the scRNA cassette. For Figure 6(E), the gRNA expression cassette was removed in its entirety from Plasmid 2.

Promoter, ribosome binding site (RBS), and terminator sequences used in these constructs are listed in Tables S2, S4, and S3, respectively, while the scRNA and gRNA sequences are provided in Table S5.

---

**Fig. S1.**
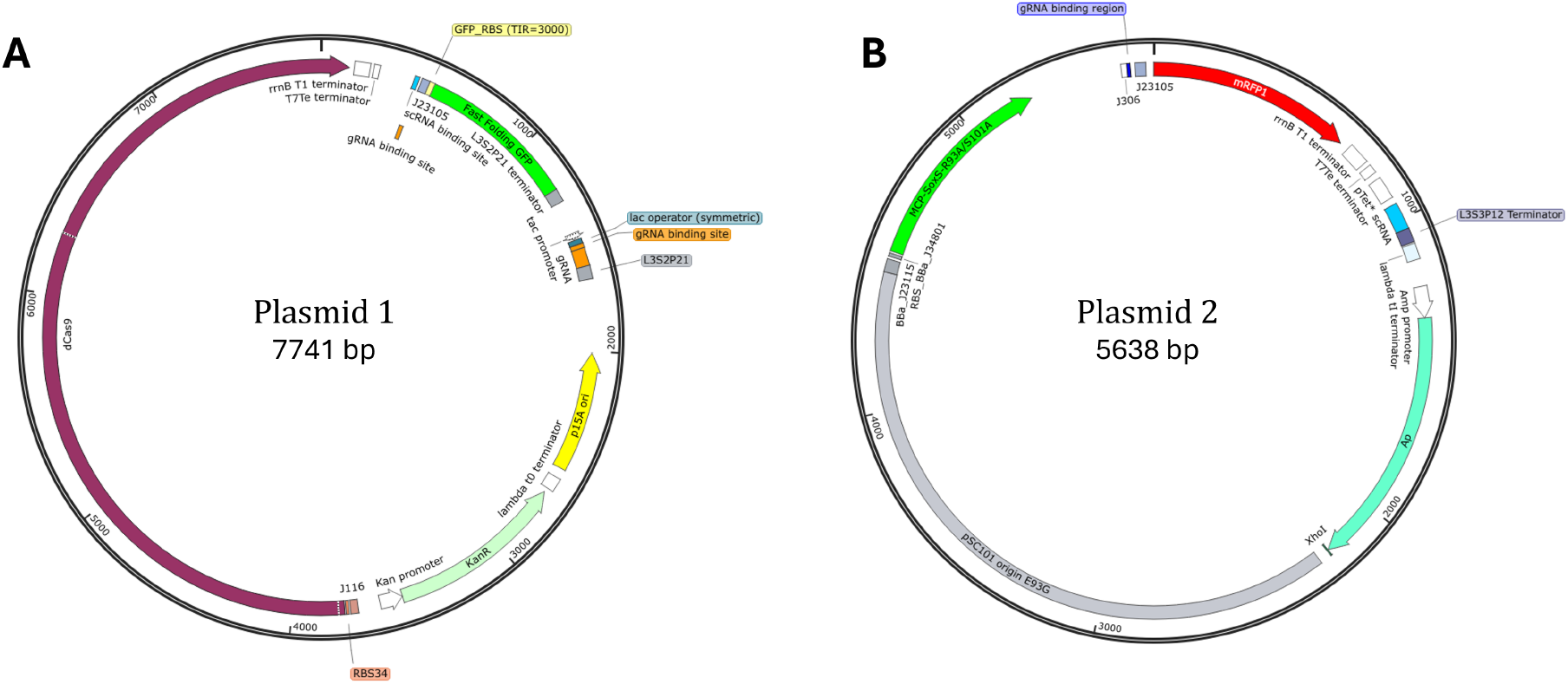
Genetic maps of plasmid constructs used in this study. (A) Genetic map of Plasmid 1, a p15A-kanamycin backbone containing constitutive dCas9 expression, a IPTG-inducible gRNA cassette, and a GFP reporter with upstream scRNA and gRNA binding sites. (B) Genetic map of Plasmid 2, a pSC101(E93G)-ampicillin backbone containing constitutive RBP–AD expression, an aTc-inducible scRNA cassette, and decoy binding sites upstream of an mRFP reporter with a dead RBS.

**Table S2.**
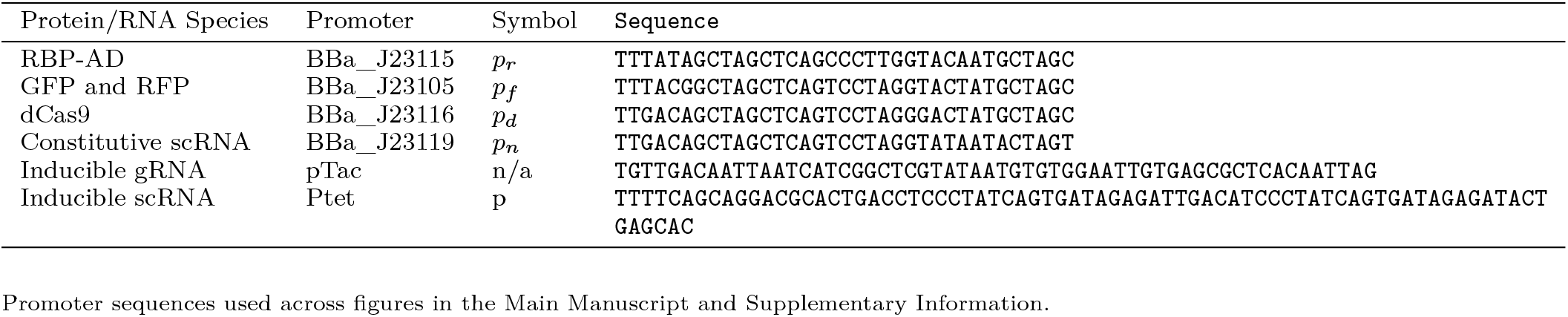
Promoter sequences used in this study.

**Table S3.**
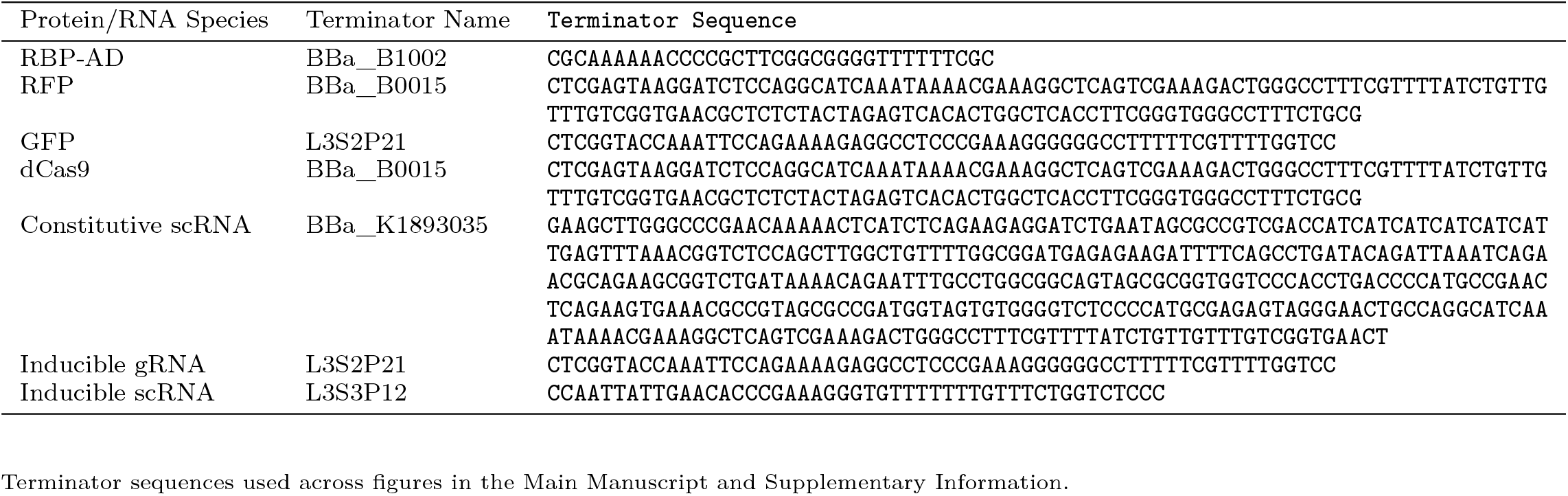
Terminator sequence used in this study.

**Table S4.**
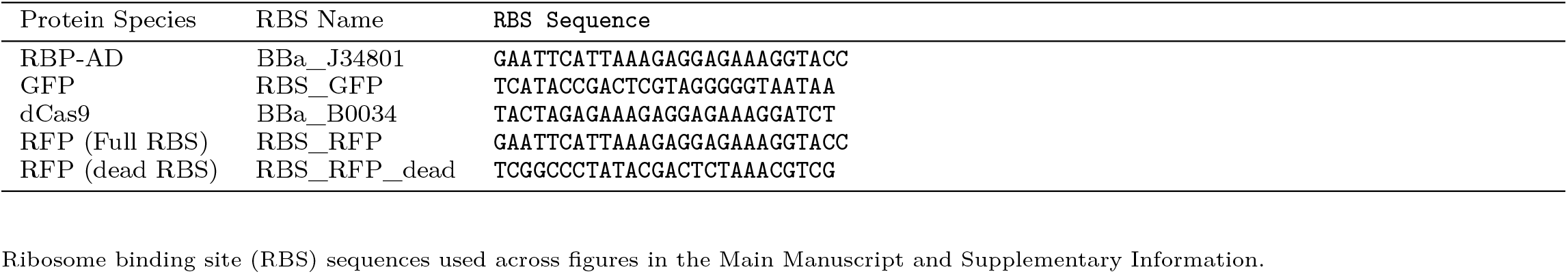
RBS sequence used in this study.

**Table S5.**
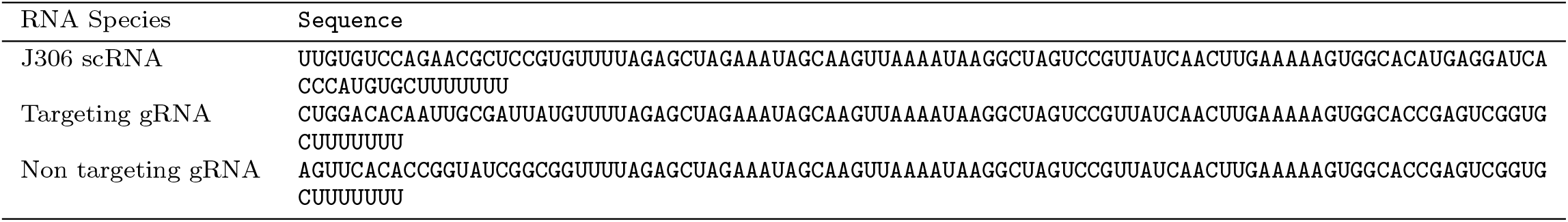
scRNA and gRNA sequences used in this study.

**Fig. S2.**
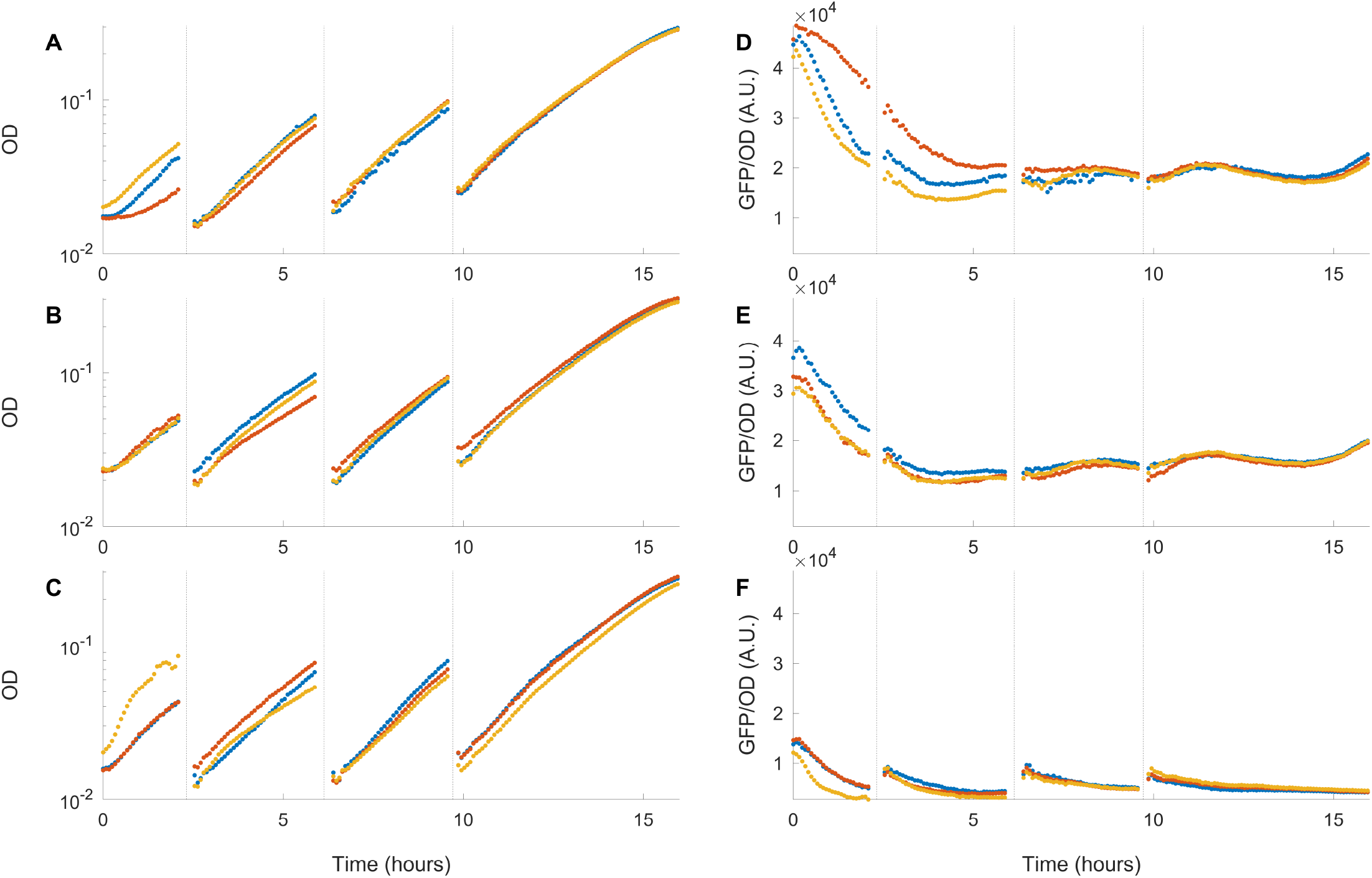
Temporal OD and GFP data corresponding to each data point presented in Main Manuscript Figure 2(C) using the genetic construct shown in Main Manuscript Figure 2(B). (A–C) Temporal OD trajectories, and (D–F, right) corresponding GFP expression over time is shown for three different conditions based on the presence or absence of scRNA and gRNA. (A,D) corresponds to {‘+ scRNA’, ‘-gRNA’}, (B,E) corresponds to the {‘−scRNA’, ‘-gRNA’}, and (C,F) corresponds to {‘−scRNA’, ‘+ gRNA’} conditions. The three colors in all panels represent three independent biological replicates. For the OD trajectories, cultures were initiated at approximately OD = 0.01 and allowed to grow to approximately OD = 0.08 before being diluted back to OD = 0.01 to maintain exponential growth. This serial dilution procedure was repeated three times, while the fourth growth cycle was allowed to proceed to stationary phase. The OD trajectory shown is representative of the temporal OD profiles observed across all experiments presented in this work. The fluorescence values reported in the Main Manuscript were taken at the time point when the OD reached 0.06 in the last batch.

**Fig. S3.**
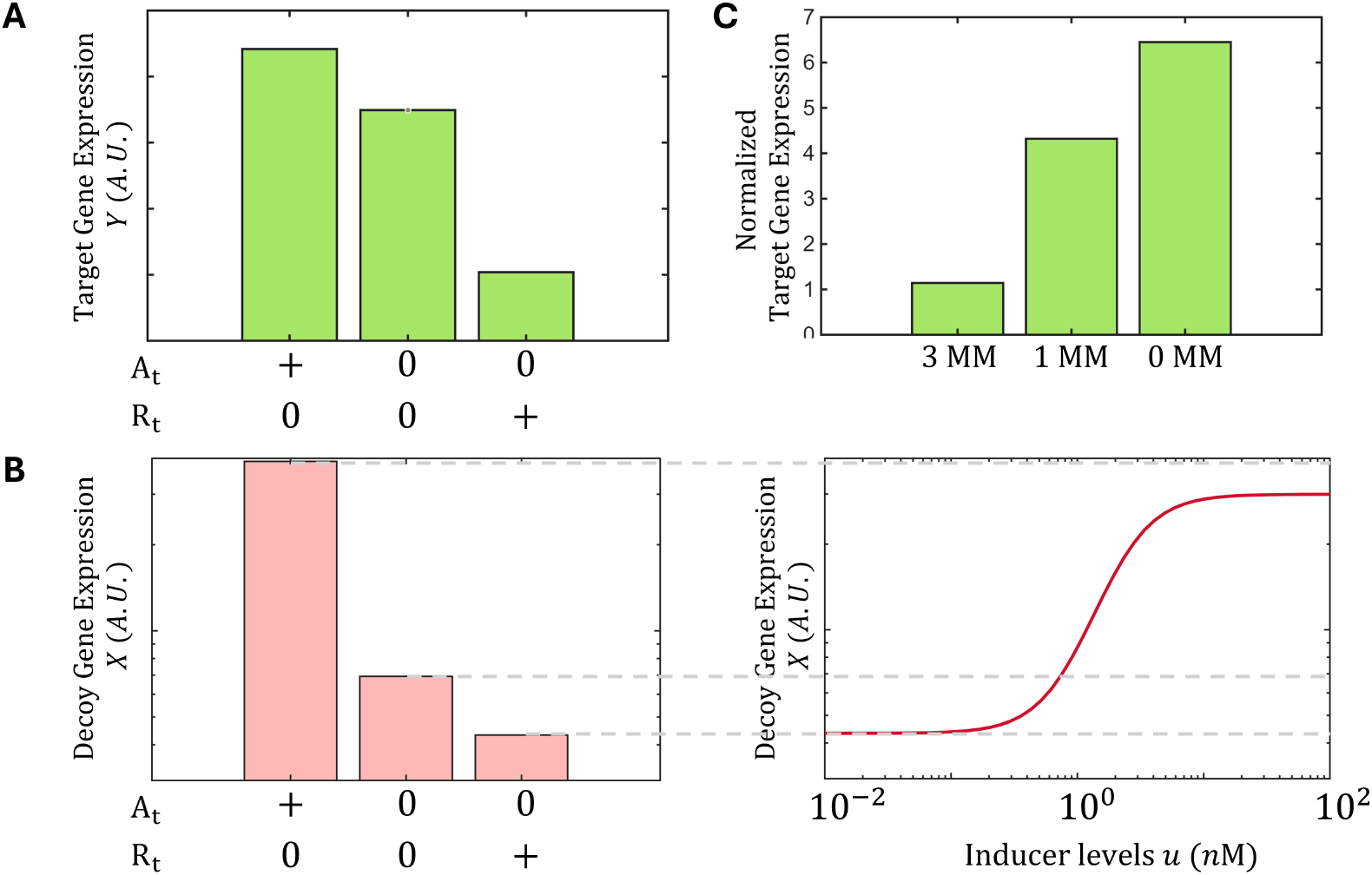
Numerical simulations of activated and repressed states for varying target binding affinities and competitive binding at the decoy site. (A) Numerical simulation of target gene expression levels (*Y*) in the activated ({*A*_*t*_, *R*_*t*_} = {+, 0}), basal ({*A*_*t*_, *R*_*t*_} = {0, 0}) and repressed ({*A*_*t*_, *R*_*t*_} = {0, +}) states of the target gene. Simulations were performed in the absence of decoy sites (i.e. *D*_*t*_ = 0) with *K*_*ta*_ = 12000. Here ‘+’ for *A*_*t*_ denotes *A*_*t*_ = 250, while ‘+’ for *R*_*t*_ denotes *R*_*t*_ = 45. Numerical simulations confirm that target gene expression is higher than the basal level in the presence of the activator and lower than the basal level in the presence of the repressor. (B) Left: Numerical simulation of decoy gene expression levels (*X*) in the activated (*{A*_*t*_, *R*_*t*_*}* = *{*+, 0*}*), basal (*{A*_*t*_, *R*_*t*_*}* = *{*0, 0*}*) and repressed (*{A*_*t*_, *R*_*t*_*}* = *{*0, +*}*) states of the decoy gene. Decoy gene expression is reduced below the basal level in the presence of the repressor alone and increased above the basal level in the presence of the activator alone. Right: Response of decoy gene expression levels (*X*) to varying activator levels induced by aTc (*u*) in the presence of the repressor (*R*_*t*_ = 45). The dashed lines indicate the extensions of the corresponding expression levels shown in the left panel. Both simulations were performed in the absence of target sites (i.e. *T*_*t*_ = 0). Increasing activator levels in the presence of a fixed amount of repressor progressively restores decoy gene expression from the repressed state to the activated state, consistent with competitive displacement of the repressor by the activator from the decoy sites. (C) Fold activation of target gene expression for different numbers of mismatches in the target–activator binding site, modeled through variation in *K*_*ta*_. old activation is defined as the ratio of *Y* at *A*_*t*_ = 250 to *Y* at *A*_*t*_ = 0. Simulations were performed in the absence of decoy sites (i.e. *D*_*t*_ = 0) for three mismatch conditions: 3 MM (*K*_*ta*_ = 12000), 1 MM (*K*_*ta*_ = 200), and 0 MM (*K*_*ta*_ = 100). Decreasing *K*_*ta*_, corresponding to stronger target–activator binding affinity, increases the fold activation levels of the target gene. The complete set of parameter values used in the simulation is provided in Table S1.

**Fig. S4.**
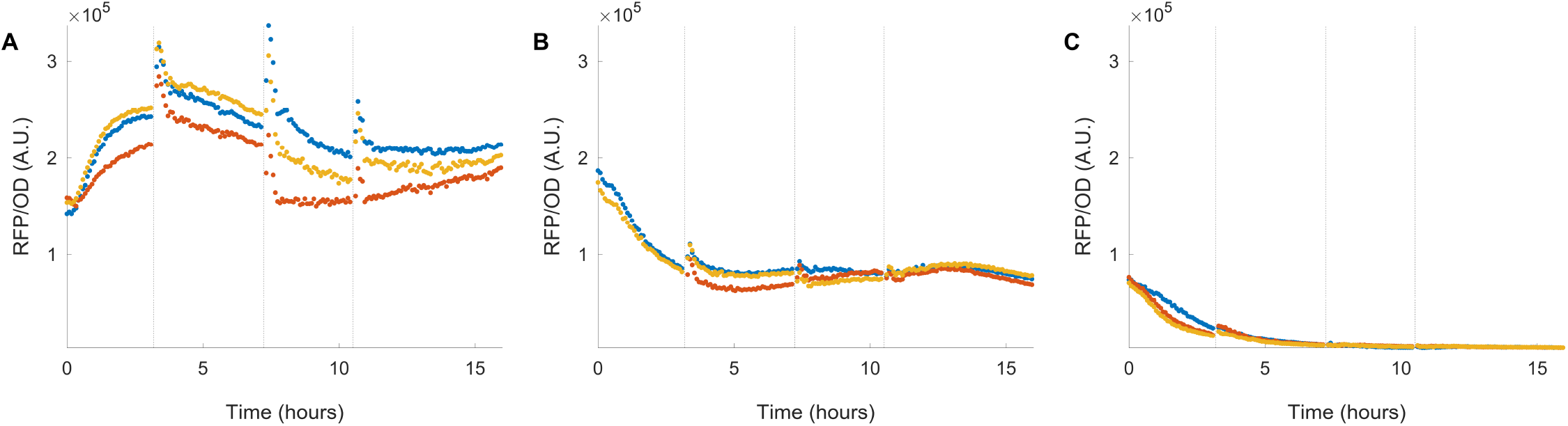
Temporal RFP data corresponding to each data point presented in Main Manuscript Figure 3(C) using the genetic construct shown in Main Manuscript Figure 3(B). Raw RFP measurements for the decoy-reporter assays summarized in Figure 3(C) with three different conditions based on the presence or absence of scRNA and gRNA. (A) corresponds to {‘+ scRNA’, ‘-gRNA’}, (B) corresponds to the {‘−scRNA’, ‘-gRNA’}, and (C) corresponds to {‘−scRNA’, ‘+ gRNA’} conditions. The three colors in all panels represent three independent biological replicates. The fluorescence values reported in the Main Manuscript were taken at the time point when the OD reached 0.06 in the last batch.

**Fig. S5.**
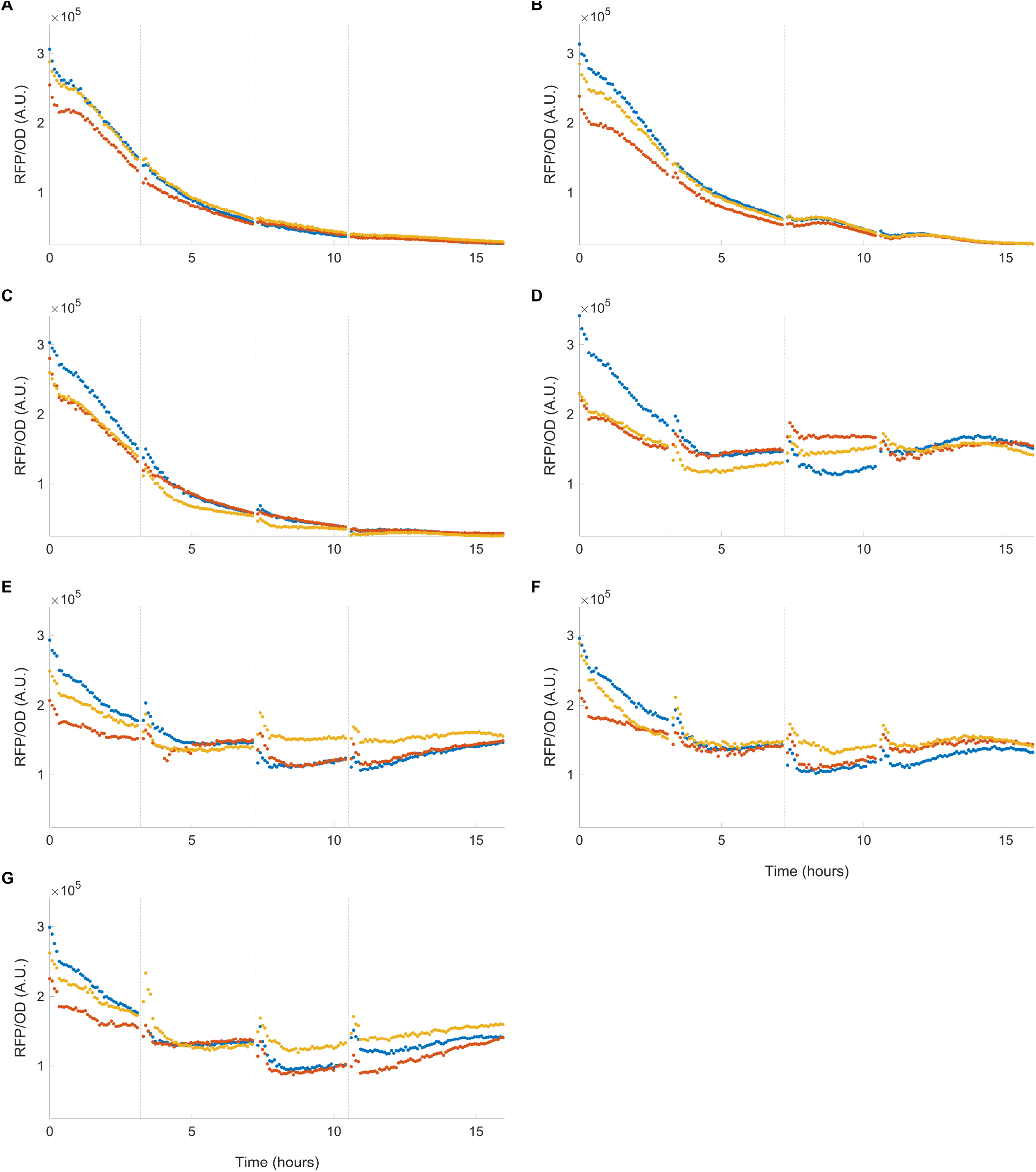
Temporal RFP data corresponding to each data point presented in Main Manuscript Figure 3(D) using the genetic construct shown in Main Manuscript Figure 3(B). (A-G) Raw RFP measurements for the decoy-reporter assays summarized for the scRNA titration experiment shown in Figure 3(D) with aTc inductions over the range [0, 0.5, 1, 3, 10, 30, 100, 300] *µ*M, respectively. The three colors in all panels represent three independent biological replicates. The fluorescence values reported in the Main Manuscript were taken at the time point when OD reached 0.06 in the last batch.

**Fig. S6.**
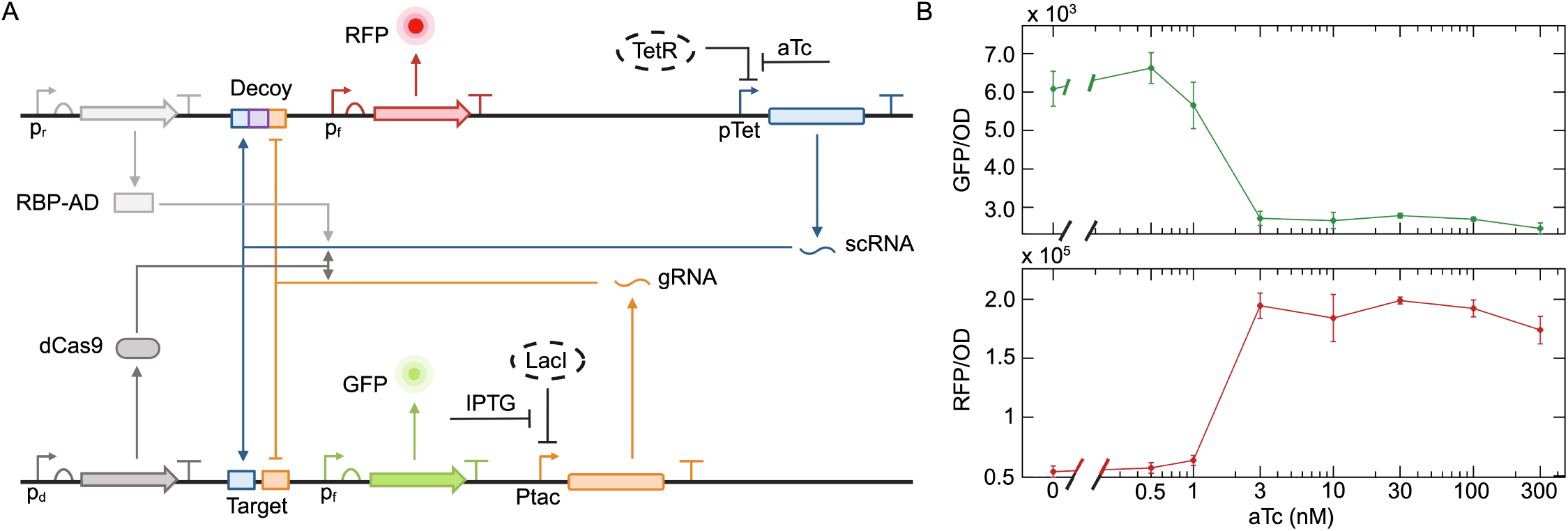
Paradoxical repression of the target gene by the activator with intact RBS for RFP enabling decoy reporter expression. (A) Genetic construct identical to that shown in Main Manuscript Figure 4(A), with the only modification being the replacement of the dead RBS downstream of the decoy sites with a functional RBS. gRNA levels are held constant using 10 *µ*M IPTG, while scRNA is induced via aTc. TetR and LacI are expressed chromosomally in the Marionette strain of E. coli (5). (B: Top) Target gene expression levels (GFP) as a function of increasing scRNA levels, induced by aTc. GFP levels decrease with increasing levels of the activator showing the emergence of paradoxical repression. (B: Bottom) RFP expression from the downstream reporter at the decoy sites. RFP levels increase with aTc, showing the binding of activators at the decoy sites. Error bars represent one standard deviation from three independent biological replicates.

**Fig. S7.**
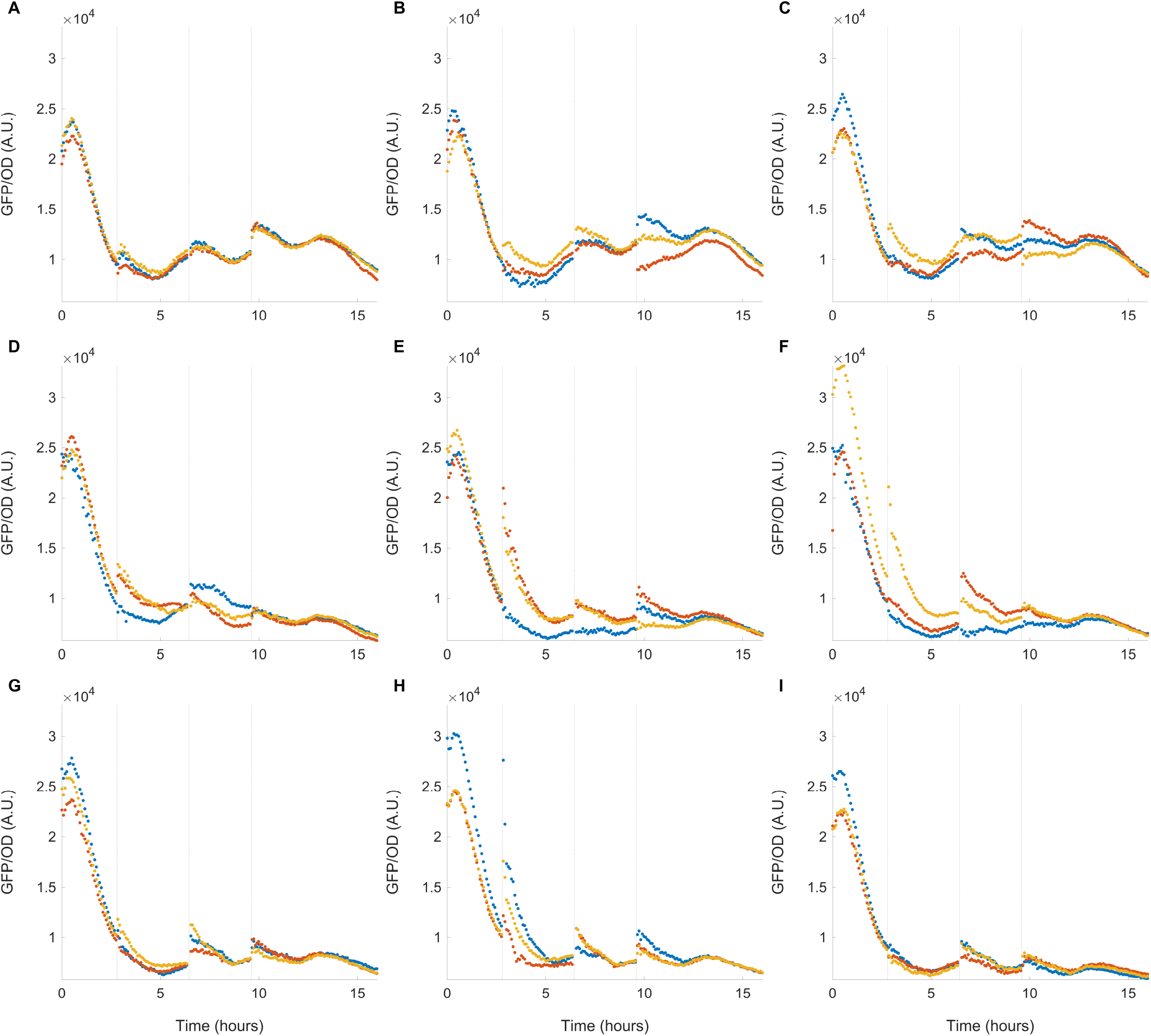
Temporal GFP data across increasing levels of activator presented in Main Manuscript Figure 4(B) using the genetic construct shown in Main Manuscript Figure 4(A). (A-H) corresponds to aTc inductions over the range [0, 0.5, 1, 3, 10, 30, 100, 300] *µ*M aTc respectively in the “-*p*_*n*_” case, i.e. without the extra constitutive scRNA cassette. (I) corresponds to 300 *µ*M aTc induction in the “+ *p*_*n*_” case, i.e. with the extra constitutive scRNA cassette. The three colors in all panels represent three independent biological replicates. The fluorescence values reported in the Main Manuscript were taken at the time point when OD reached 0.06 in the last batch. This point is normalized by dividing by the mean expression value at 0 induction, whose time series is shown in panel (A).

**Fig. S8.**
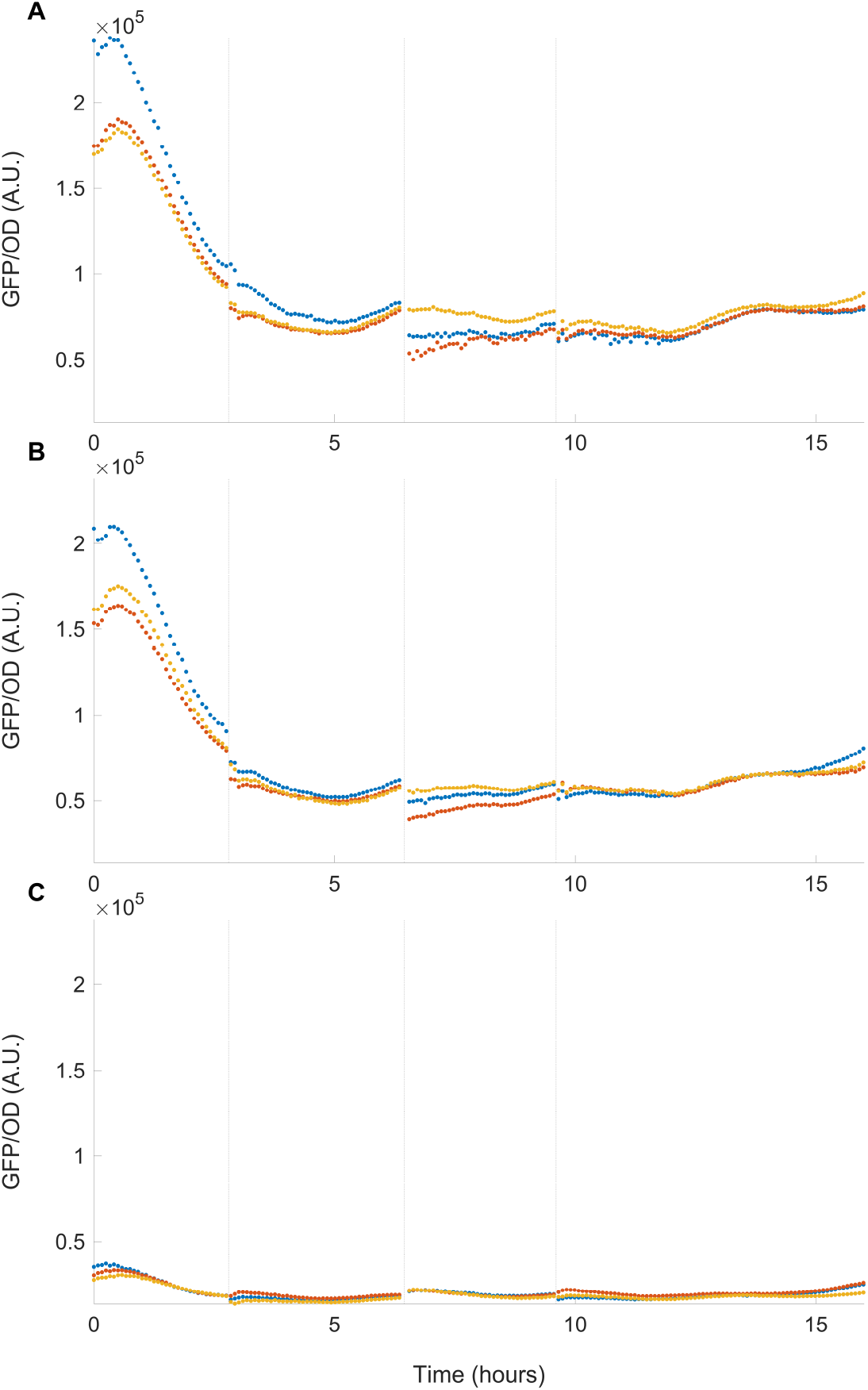
Temporal GFP data across various mismatches presented in Main Manuscript Figure 5(B) using the genetic construct shown in Main Manuscript Figure 5(B). GFP expression over time is shown for various mismatches at the target site, specifically (A) 0 MM, (B) 1 MM, and (C) 3 MM. The three colors in all panels represent three independent biological replicates. The fluorescence values reported in the Main Manuscript were taken at the time point when OD reached 0.06 in the last batch. This point is normalized by dividing by the mean expression value at 0 induction, whose time series is shown in panel (A).

**Fig. S9.**
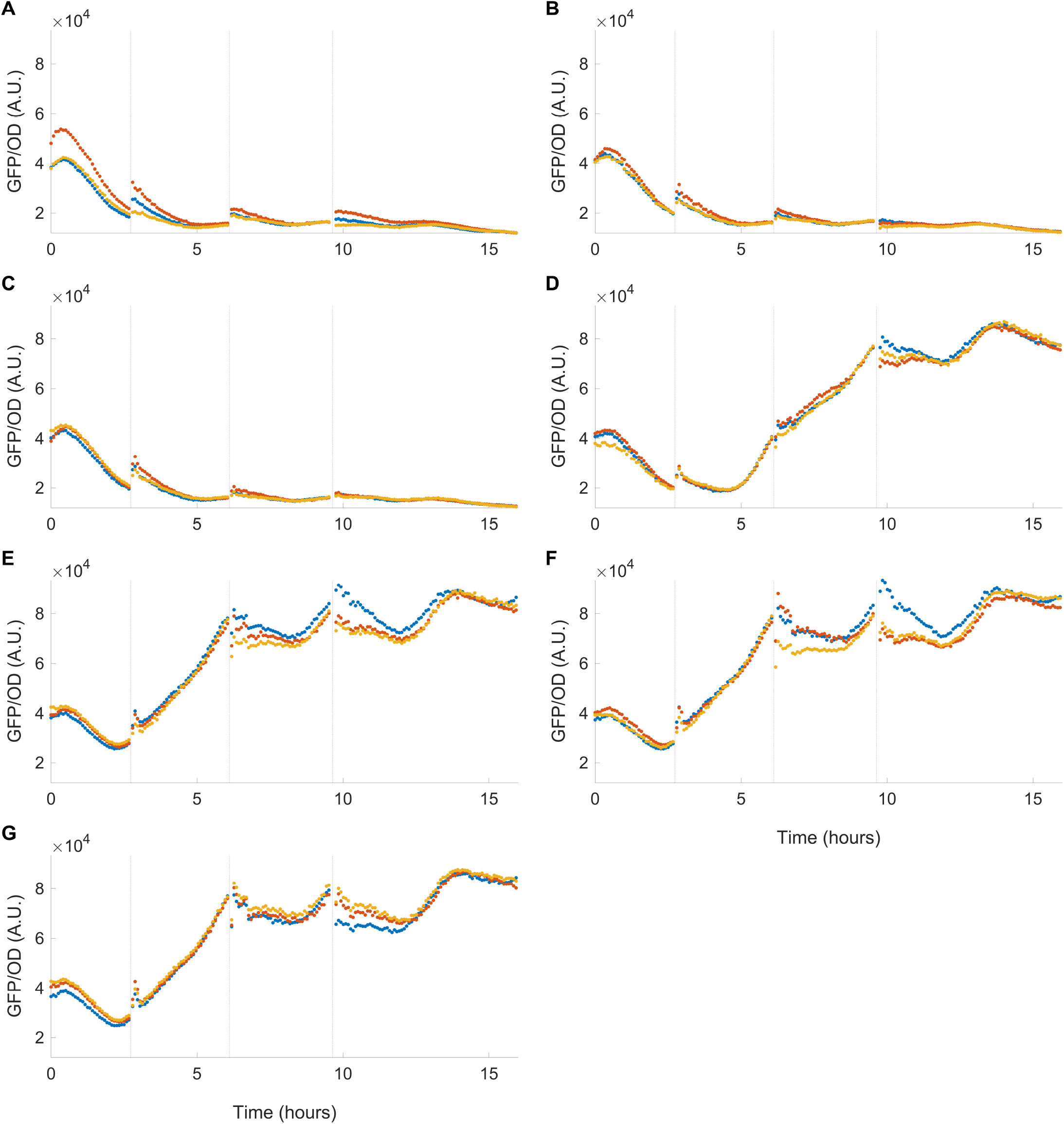
Temporal GFP data across increasing levels of activator presented in Main Manuscript Figure 5(D) using the genetic construct shown in Main Manuscript Figure 5(C) with 0 MM at the target site. (A-G) GFP expression over time for aTc inductions over the range of [0, 0.5, 1, 3, 10, 30, 100] *µ*M, respectively. The three colors in all panels represent three independent biological replicates. The fluorescence values reported in the Main Manuscript were taken at the time point when OD reached 0.06 in the last batch. This point is normalized by dividing by the mean expression value at 0 induction, whose time series is shown in panel (A).

**Fig. S10.**
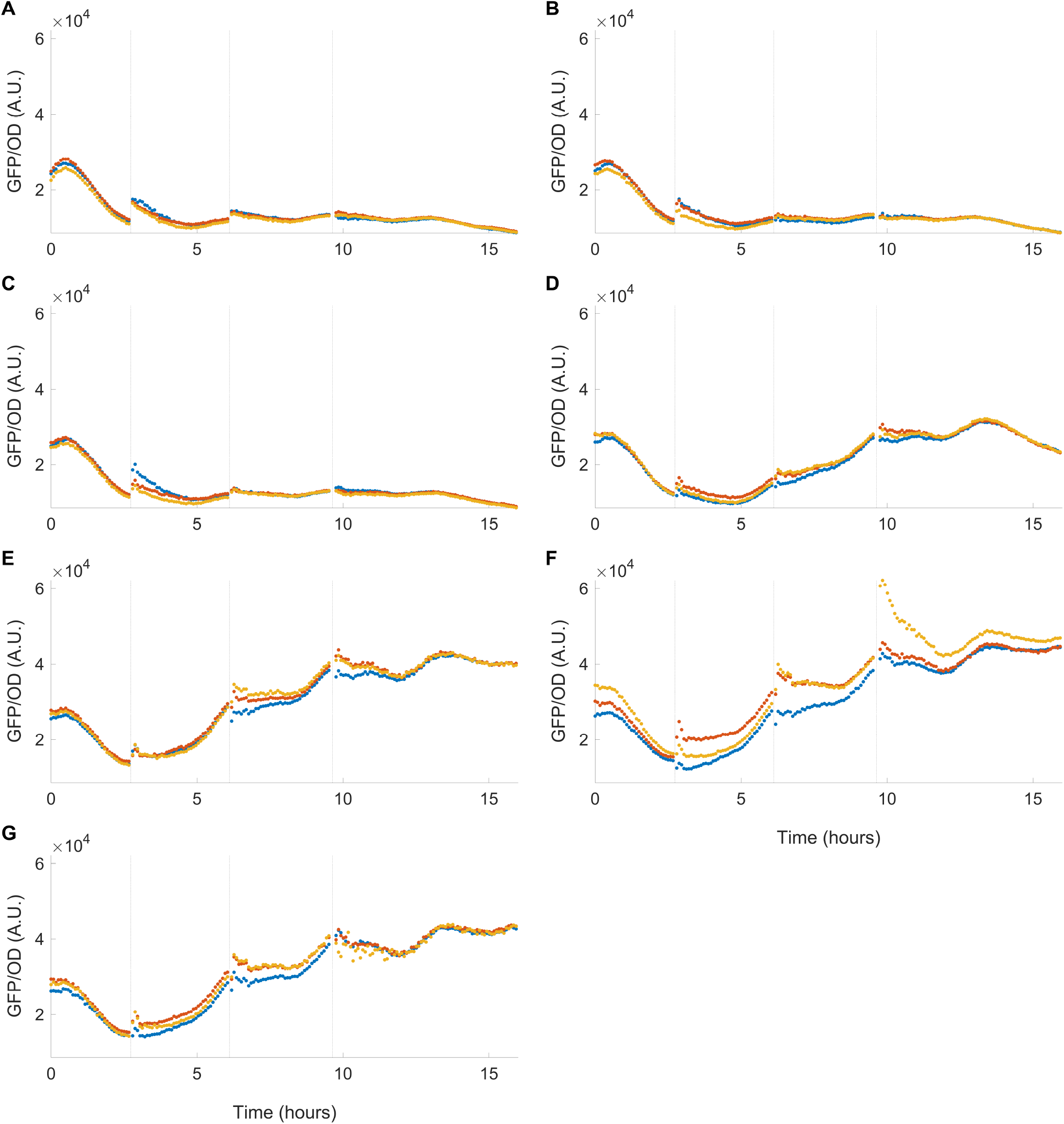
Temporal GFP data across increasing levels of activator presented in Main Manuscript Figure 5(D) using the genetic construct shown in Main Manuscript Figure 5(C) with 1 MM at the target site. (A-G) GFP expression over time for aTc inductions over the range of [0, 0.5, 1, 3, 10, 30, 100] *µ*M, respectively. The three colors in all panels represent three independent biological replicates. The fluorescence values reported in the Main Manuscript were taken at the time point when OD reached 0.06 in the last batch. This point is normalized by dividing by the mean expression value at 0 induction, whose time series is shown in panel (A).

**Fig. S11.**
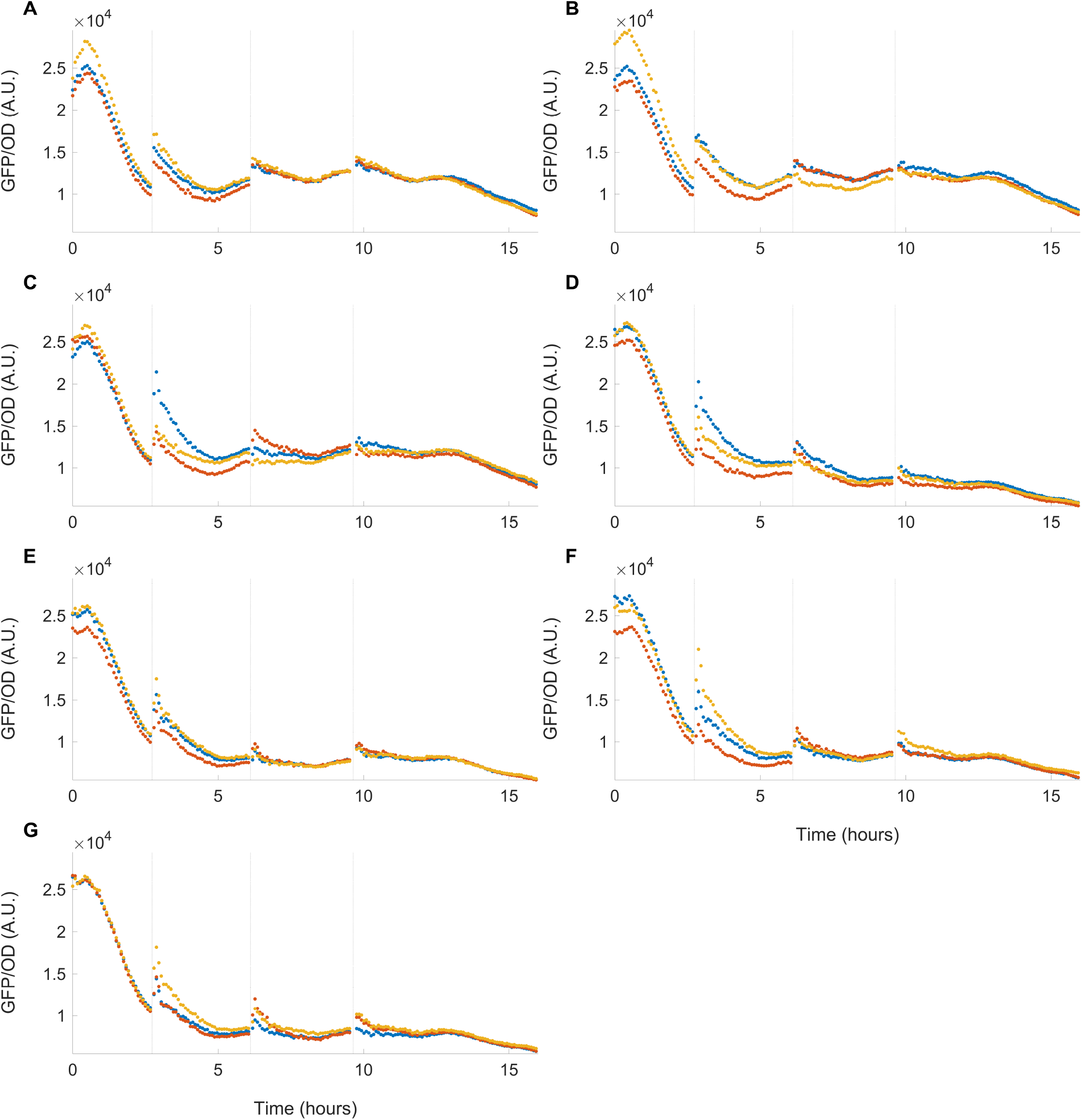
Temporal GFP data across increasing levels of activator presented in Main Manuscript Figure 5(D) using the genetic construct shown in Main Manuscript Figure 5(C) with 3 MM at the target site. (A-G) GFP expression over time for aTc inductions over the range of [0, 0.5, 1, 3, 10, 30, 100] *µ*M, respectively. The three colors in all panels represent three independent biological replicates. The fluorescence values reported in the Main Manuscript were taken at the time point when OD reached 0.06 in the last batch. This point is normalized by dividing by the mean expression value at 0 induction, whose time series is shown in panel (A).

**Fig. S12.**
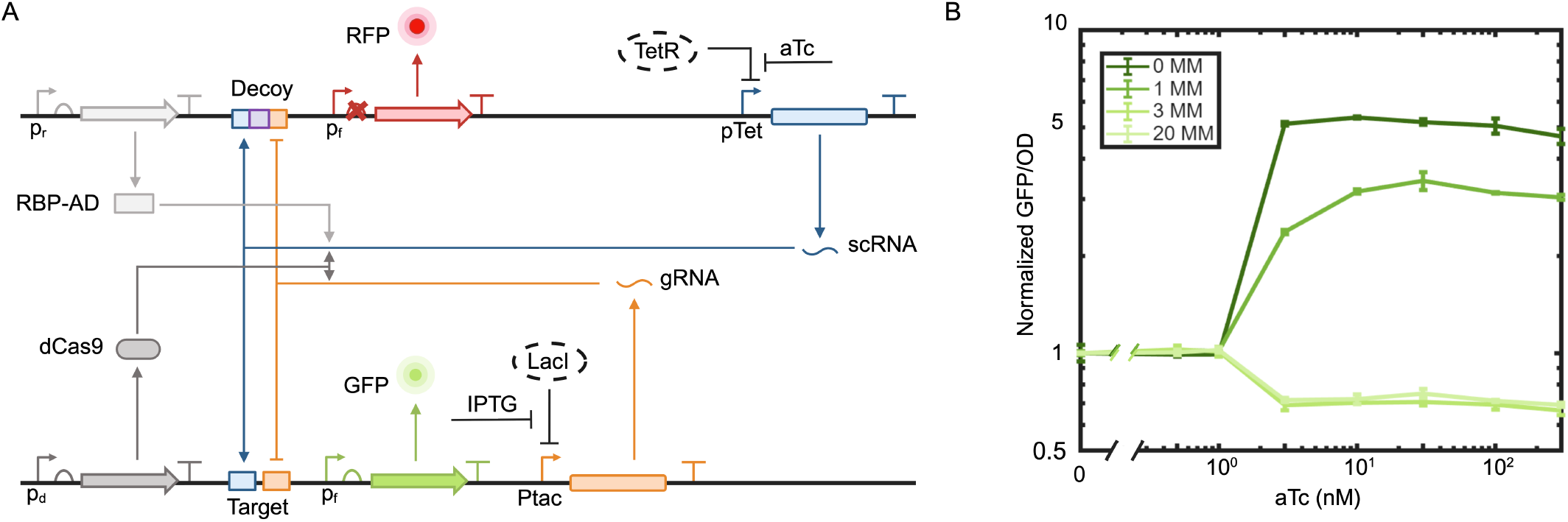
Effect of target–activator binding affinity on the emergence of paradoxical repression. (A) Genetic circuit identical to that shown in Main Manuscript Figure 5(C), with constant repressor expression, overlapping binding at decoy sites, and variable scRNA levels via aTc induction. (B) Normalized GFP output as a function of increasing scRNA levels for different target–activator binding affinities, tuned by varying the number of mismatches between the scRNA and target site as 0 (0 MM), 1 (1 MM), 3 (3 MM), and 20 (20 MM) mismatches. GFP values are normalized with the corresponding GFP levels at 0 aTc (no induction). The 20 MM condition corresponds to essentially no binding between activator and target and exhibits paradoxical repression behavior. In contrast, reducing mismatches progressively increases binding affinity, with 3 MM showing weak activation and paradoxical repression, 1 MM and 0 MM showing the stronger activation and complete loss of paradoxical behavior. These results demonstrate that weak target–activator binding is essential to observe paradoxical repression with increasing scRNA levels.

**Fig. S13.**
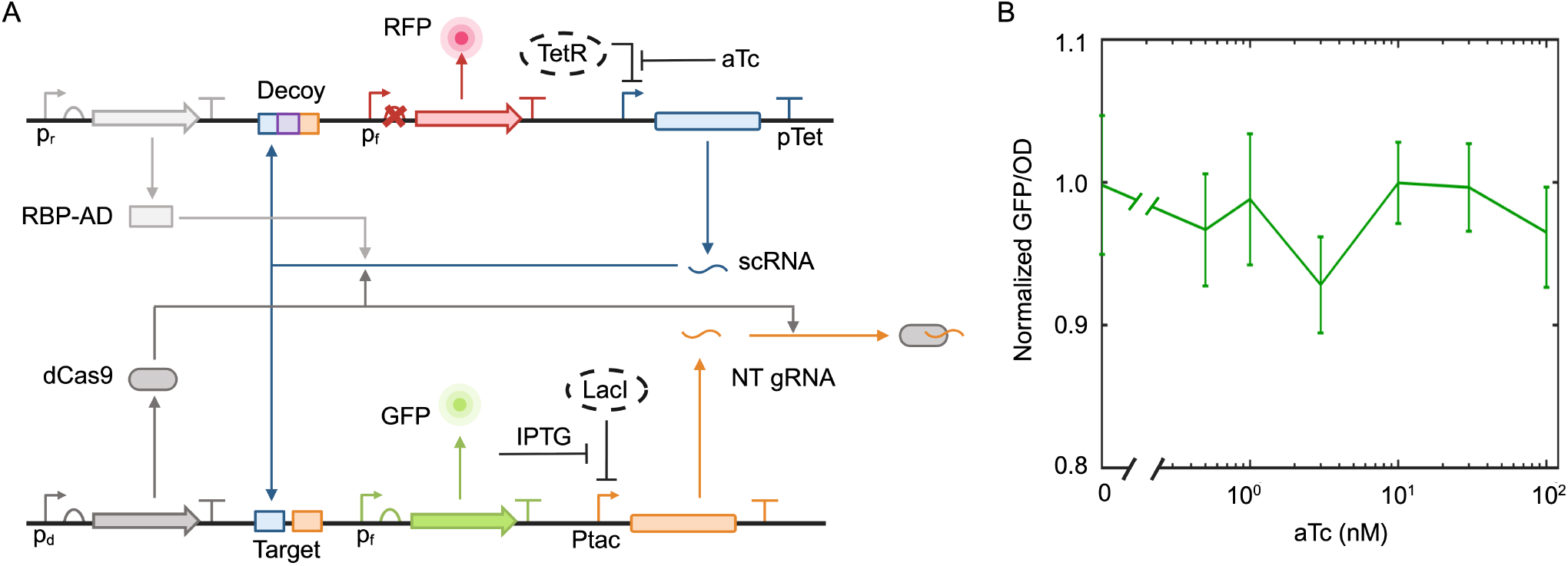
Paradoxical repression disappears in the presence of non-target gRNA. (A) Genetic circuit identical to that in Main Manuscript Figure 5(C) with 3MM mismatch at the target site, except that a non-target gRNA (NT gRNA) is used instead of the target gRNA. The NT gRNA does not bind either the target sequence or the decoy sites. However, it still binds dCas9 to form a dCas9–NT-gRNA complex, serving as a control for dCas9 competition effects. (B) Normalized GFP output as a function of increasing scRNA levels in the presence of NT gRNA. GFP values are normalized with the GFP level at 0 aTc (no induction). Unlike with target gRNA, paradoxical repression is no longer observed, and target expression remains close to basal levels.

**Fig. S14.**
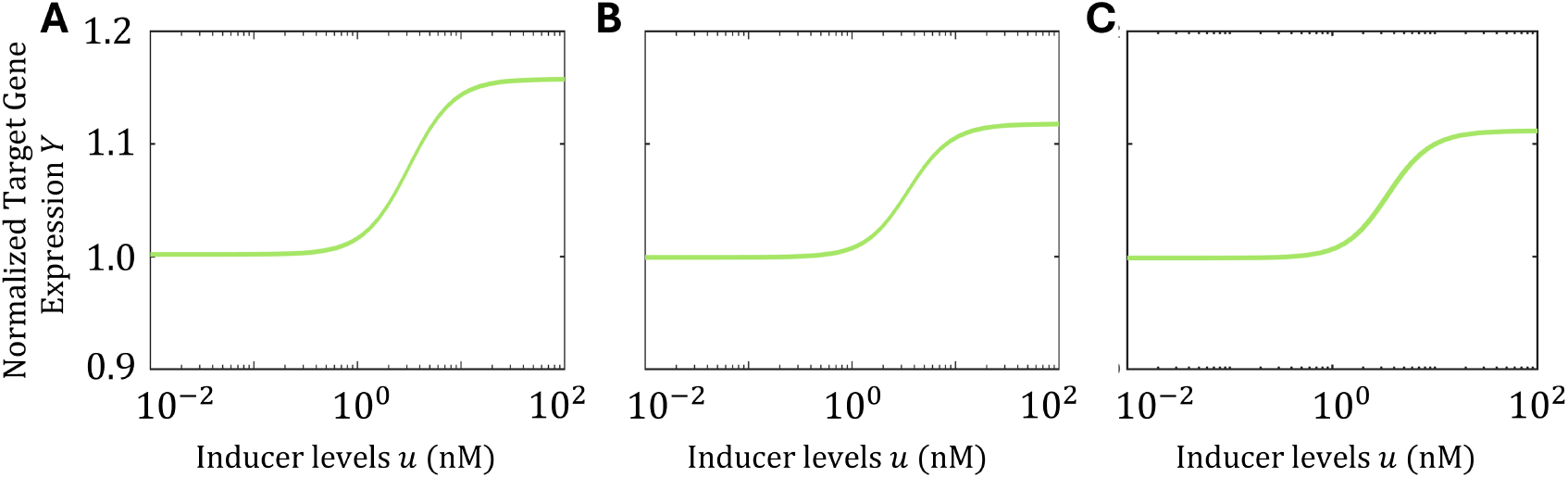
Numerical figure showing the absence of paradoxical gene expression when critical requirements are not met. (A) Input–output response of normalized target gene expression *Y* to increasing activator levels via aTc induction *u* in the absence of decoy sites (*D*_*t*_ = 0), corresponding to Figure 6(A–B) in the main manuscript. The system no longer exhibits paradoxical repression and instead shows a slight activation. (B) Input output response of *Y* for increasing *u* in a system without overlapping decoy sites, where activator and repressor bind to separate decoys (*D*_1_ and *D*_2_) forming *D*_1*A*_ and *D*_2*R*_, respectively. The total concentrations are *D*_*t*_ = *D*_1_ + *D*_1*A*_ = *D*_2_ + *D*_2*R*_, *A*_*t*_ = *A* + *T*_*A*_ + *D*_1*A*_, *R*_*t*_ = *R* + *T*_*R*_ + *D*_2*R*_, and *T*_*t*_ = *T* + *T*_*A*_ + *T*_*R*_. This corresponds to Figure 6(C–D) in the main manuscript and also shows loss of paradoxical repression with slight activation. (C) Input output response of *Y* for increasing *u* in the absence of repressor (*R*_*t*_ = 0), corresponding to Figure 6(E–F) in the main manuscript, where the system again shows no paradoxical repression and instead a slight activation. The target expression values (*Y*) in (A), (B) and (C) are normalized with the corresponding expression values at no induction, *Y* (*u* = 0 nM). The complete set of parameter values used in the simulation is provided in Table S1.

**Fig. S15.**
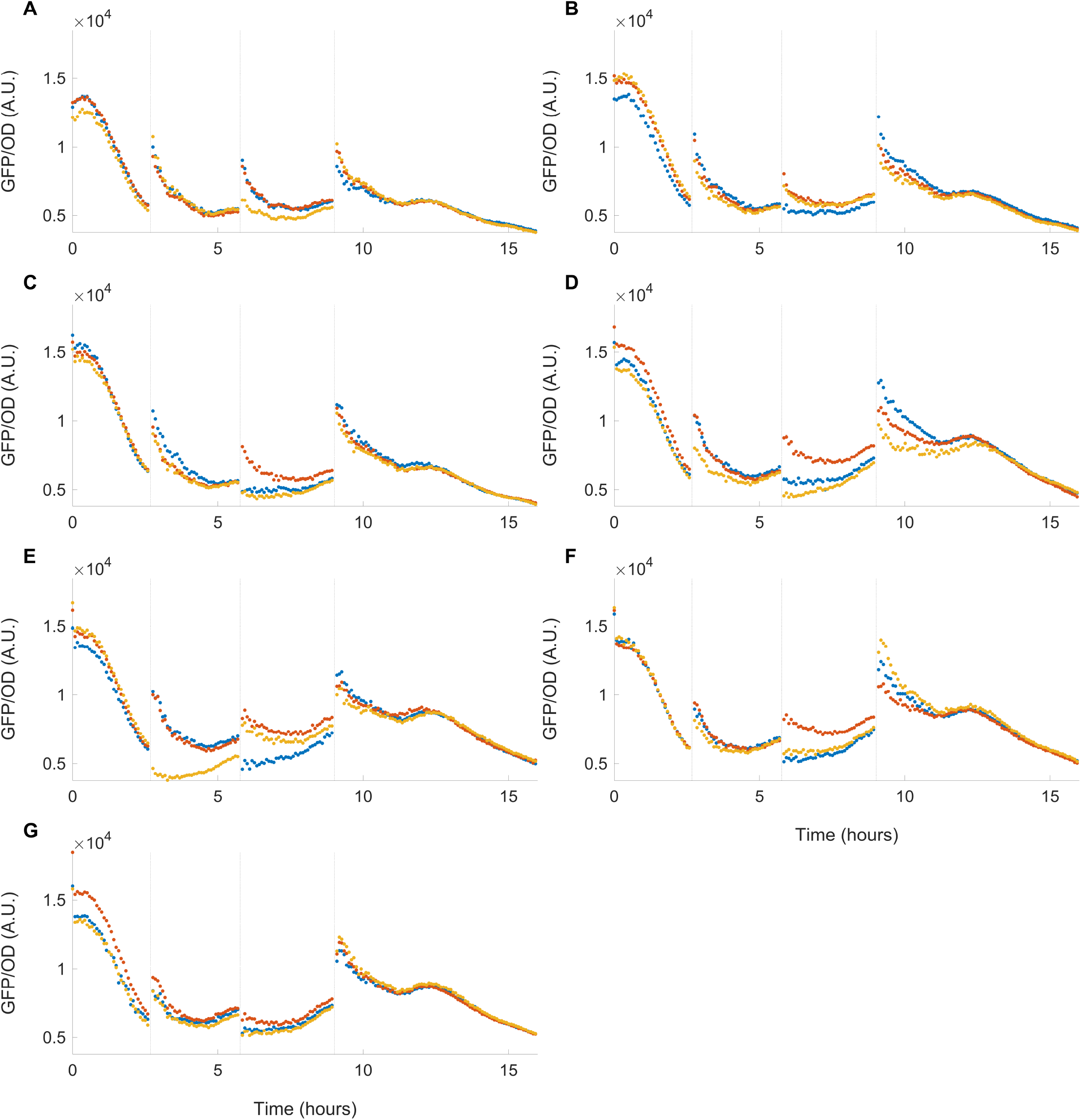
Temporal GFP data across increasing levels of activator presented in Main Manuscript Figure 6(B) using the genetic construct shown in Main Manuscript Figure 6(A). (A-G) GFP expression over time for aTc inductions over the range of [0, 0.5, 1, 3, 10, 30, 100] *µ*M, respectively with all critical system components except for the decoy, and with 3MM at the target site. The three colors in all panels represent three independent biological replicates. The fluorescence values reported in the Main Manuscript were taken at the time point when OD reached 0.06 in the last batch. This point is normalized by dividing by the mean expression value at 0 induction, whose time series is shown in panel (A).

**Fig. S16.**
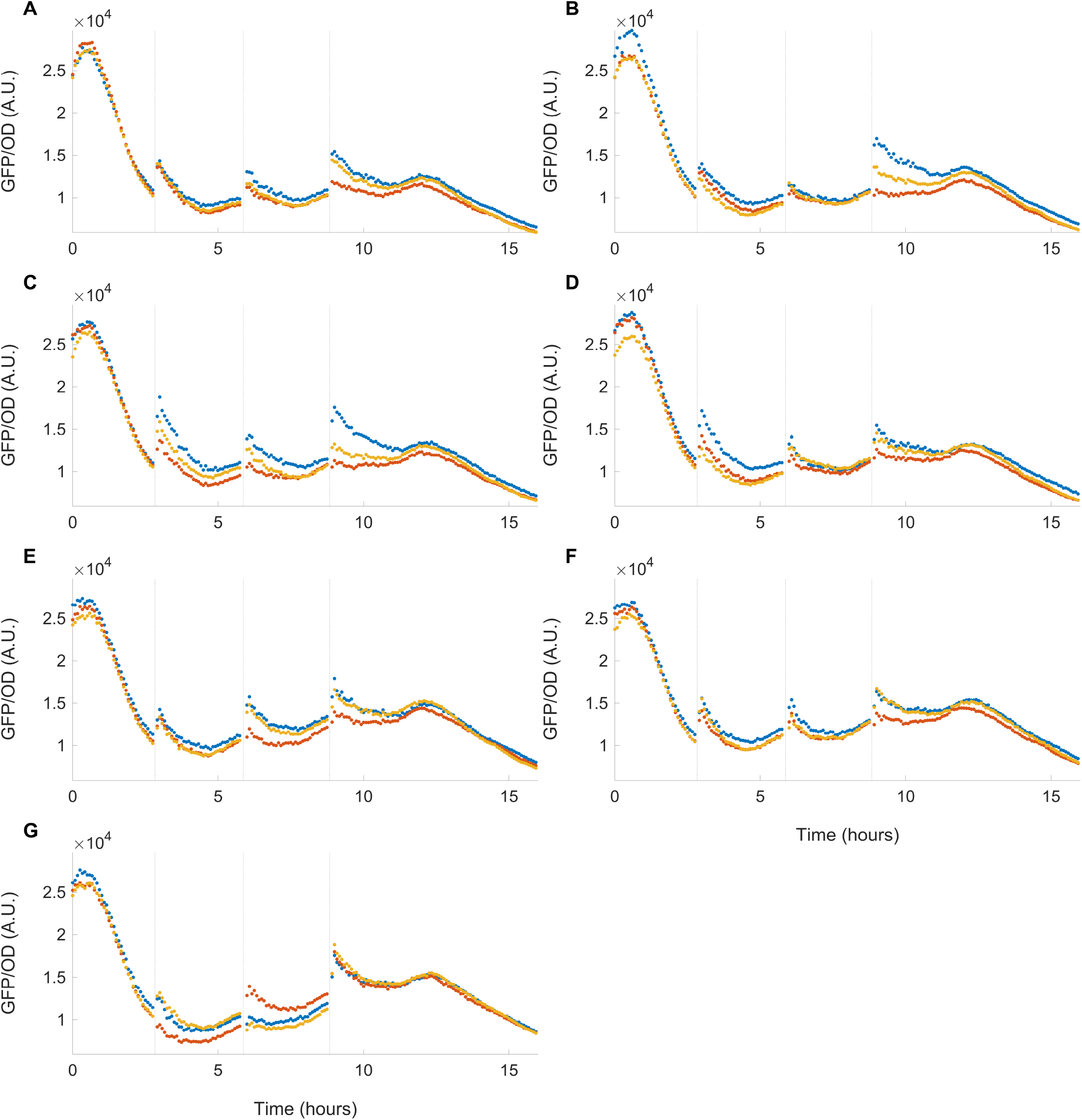
Temporal GFP data across increasing levels of activator presented in Main Manuscript Figure 6(D) using the genetic construct shown in Main Manuscript Figure 6(C). (A-G) GFP expression over time for aTc inductions over the range of [0, 0.5, 1, 3, 10, 30, 100] *µ*M, respectively with non-overlapping binding sites at the decoy, and with 3MM at the target site. The three colors in all panels represent three independent biological replicates. The fluorescence values reported in the Main Manuscript were taken at the time point when OD reached 0.06 in the last batch. This point is normalized by dividing by the mean expression value at 0 induction, whose time series is shown in panel (A).

**Fig. S17.**
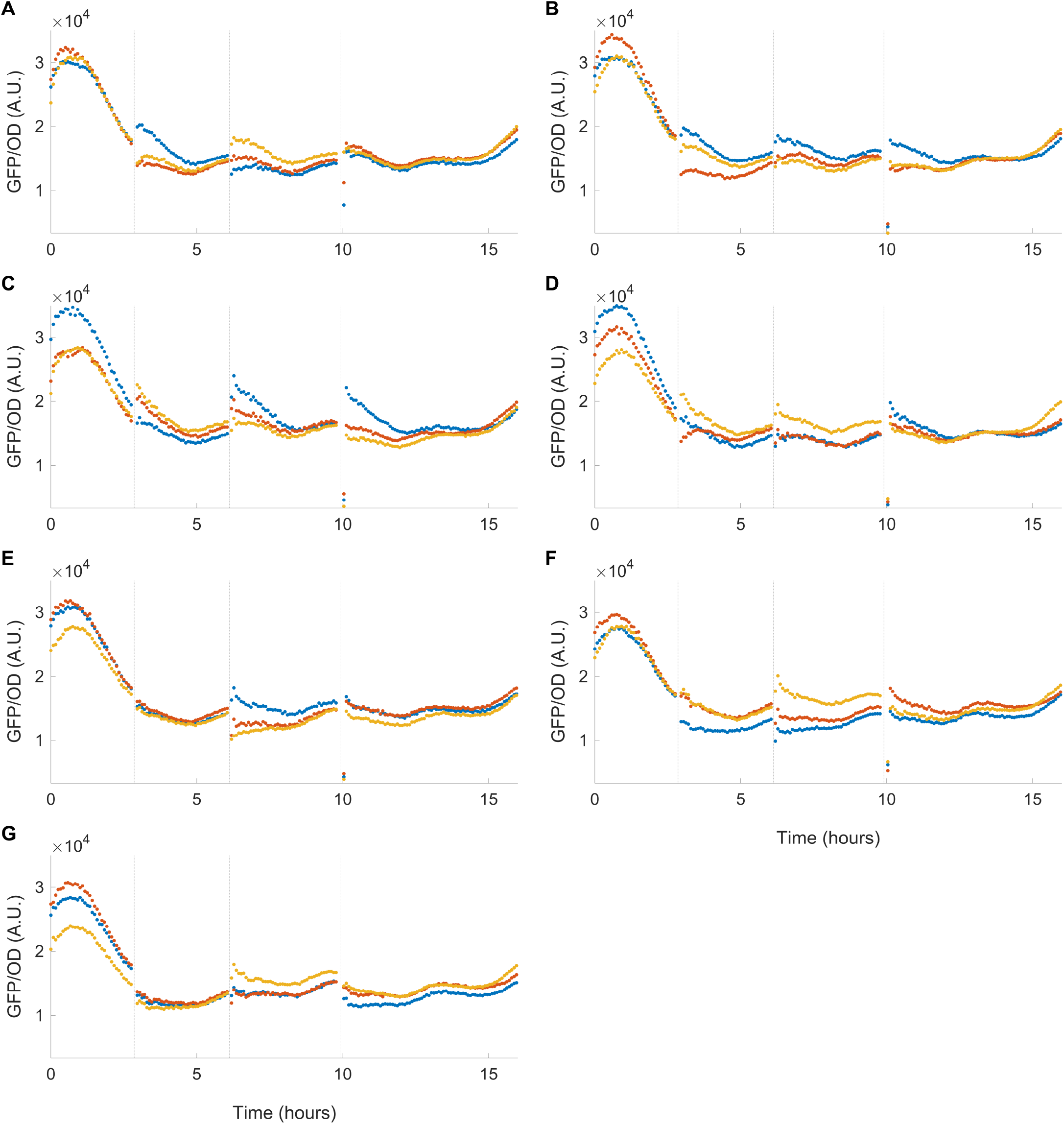
Temporal GFP data across increasing levels of activator presented in Main Manuscript Figure 6(F) using the genetic construct shown in Main Manuscript Figure 6(E). (A-G) GFP expression over time for aTc inductions over the range of [0, 0.5, 1, 3, 10, 30, 100] *µ*M, respectively with all critical system components except for the repressor, and with 3MM at the target site. The three colors in all panels represent three independent biological replicates. The fluorescence values reported in the Main Manuscript were taken at the time point when OD reached 0.06 in the last batch. This point is normalized by dividing by the mean expression value at 0 induction, whose time series is shown in panel (A).

**Fig. S18.**
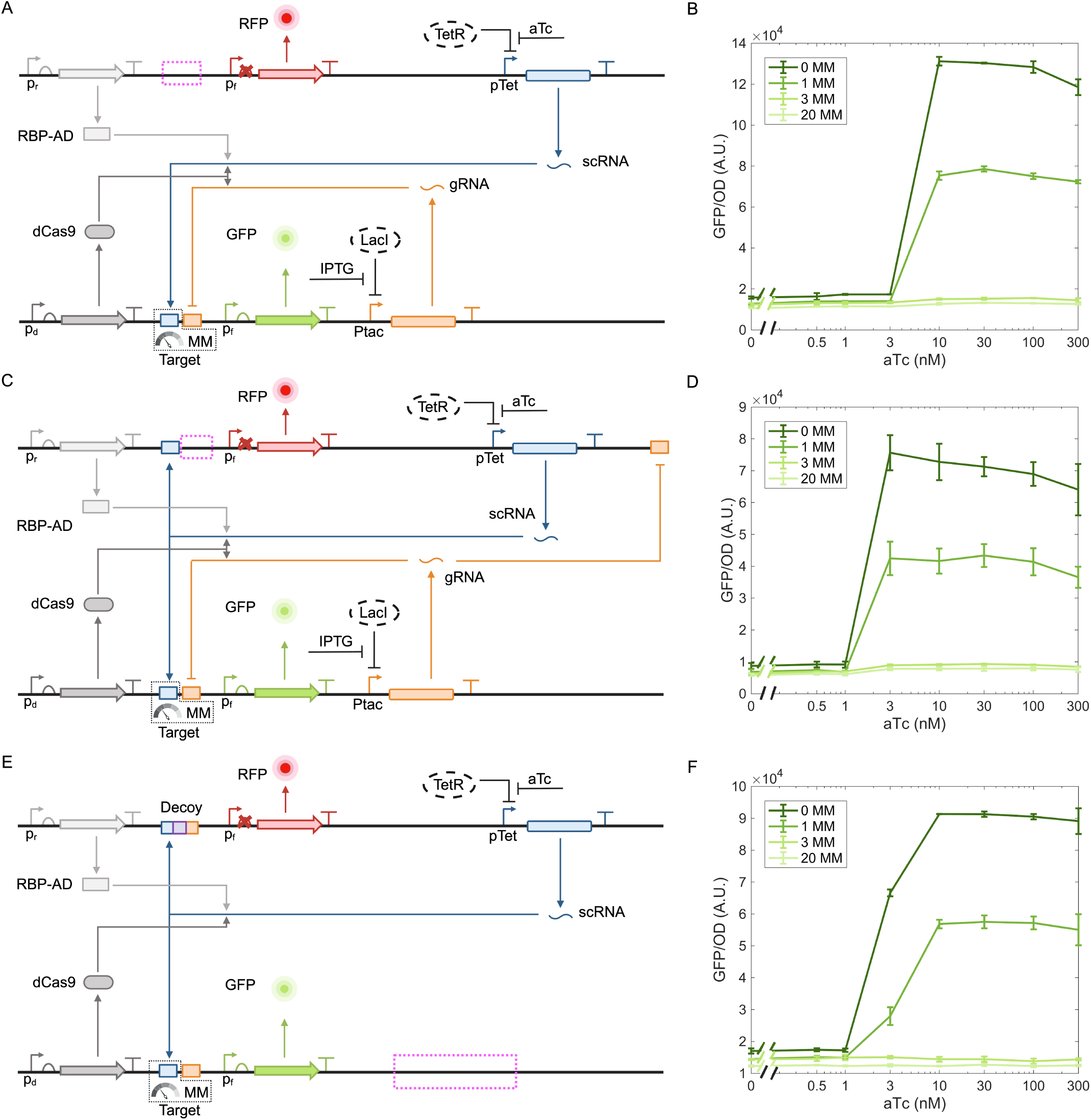
Effect of target-site mismatches on the paradoxical response in constructs lacking decoy sites, overlapping binding, or repressor binding. (A) Genetic construct lacking decoy sites with target sites containing 0, 1, 3, or 20 mismatches (MM). (B) GFP input–output response to increasing aTc induction for the constructs shown in (A) under 10 *µ*M IPTG induction. Removing the decoy sites from the genetic construct eliminated the paradoxical regulation of 3 MM and 20 MM, causing GFP expression to increase with higher scRNA levels. For the 0 MM and 1 MM target-site conditions, increasing scRNA levels led to increased GFP expression, resulting in higher output levels upon activation. (C) Genetic construct lacking overlapping competitive binding at the decoy sites with target sites containing 0, 1, 3, or 20 MM. (D) GFP input–output response to increasing aTc induction for the constructs shown in (C) under 10 *µ*M IPTG induction. In this configuration, GFP expression increased with scRNA levels for 3 MM and 20 MM, indicating the absence of paradoxical regulation. For both the 0 MM and 1 MM target-site conditions, no significant differences in behavior were observed. Overall expression was lower than in the no-decoy case due to sequestration of activators by the non-overlapping decoy sites. (E) Genetic construct lacking repressor binding with target sites containing 0, 1, 3, or 20 MM. (F) GFP input–output response to increasing aTc induction for the constructs shown in (E). GFP expression remained largely constant across increasing aTc concentrations for 3 MM and 20 MM, indicating a loss of the paradoxical response. For the 0 MM and 1 MM conditions, a slight increase in expression is observed compared to the repressor-present case, likely due to residual sequestration of activators by the decoy sites. In all constructs, plasmids were transformed into the Marionette strain of E. coli, where the corresponding regulators are expressed endogenously. Error bars represent one standard deviation from three independent biological replicates. Fluorescence values are reported in arbitrary units (A.U.).

